# Modelling the effects of environmental heterogeneity within the lung on the tuberculosis life-cycle

**DOI:** 10.1101/2019.12.12.871269

**Authors:** Michael J. Pitcher, Ruth Bowness, Simon Dobson, Raluca Eftimie, Stephen H. Gillespie

## Abstract

Progress in shortening the duration of tuberculosis (TB) treatment is hampered by the lack of a predictive model that accurately reflects the diverse environment within the lung. This is important as TB has been shown to produce distinct localisations to different areas of the lung during different disease stages, with the environmental heterogeneity within the lung of factors such as air ventilation, blood perfusion and oxygen tension believed to contribute to the apical localisation witnessed during the post-primary form of the disease.

Building upon our previous model of environmental lung heterogeneity, we present a networked metapopulation model that simulates TB across the whole lung, incorporating these notions of environmental heterogeneity across the whole TB life-cycle to show how different stages of the disease are influenced by different environmental and immunological factors. The alveolar tissue in the lung is divided into distinct patches, with each patch representing a portion of the total tissue and containing environmental attributes that reflect the internal conditions at that location. We include populations of bacteria and immune cells in various states, and events are included which determine how the members of the model interact with each other and the environment. By allowing some of these events to be dependent on environmental attributes, we create a set of heterogeneous dynamics, whereby the location of the tissue within the lung determines the disease pathological events that occur there.

Our results show that the environmental heterogeneity within the lung is a plausible driving force behind the apical localisation during post-primary disease. After initial infection, bacterial levels will grow in the initial infection location at the base of the lung until an adaptive immune response is initiated. During this period, bacteria are able to disseminate and create new lesions throughout the lung. During the latent stage, the lesions that are situated towards the apex are the largest in size, and once a post-primary immune-suppressing event occurs, it is the uppermost lesions that reach the highest levels of bacterial proliferation. Our sensitivity analysis also shows that it is the differential in blood perfusion, causing reduced immune activity towards the apex, which has the biggest influence of disease outputs.

## 1. Introduction

Tuberculosis (TB) accounts for over 1 million deaths each year [1], despite the fact that an effective treatment has existed for decades. The current standard regimen for drug-susceptible forms of TB requires six months of multiple antibiotics, and a large number of factors can contribute to a patient’s ability to adhere to the treatment [2]. Non-adherence can have serious consequences, both for the patient, as it increases the chances of relapse after treatment, and for the wider society, as an incomplete course of antibiotics can lead to the remaining bacteria developing drug resistance [3, 4]. Therefore, creating novel regimens of shorter duration is of great importance, as doing so would improve overall patient adherence and reduce these risks of relapse and drug resistance. Unfortunately, recent efforts to create new regimens of four months have not been successful.

Experiments on mice using moxifloxacin showed promising results with regard to bactericidal effects [5] and it was predicted that the use of this drug could reduce human treatment duration. However, clinical trials incorporating the drug in novel regimens were unable to prove non-inferiority [6, 7], possibly due to the heterogeneity of distribution of drugs within the lesion during treatment [8]. This demonstrates a crucial hurdle in the drug development process for TB: we lack the predictive power at the preclinical stage to make effective decisions as to which of the many possible new regimens to progress through to expensive and costly clinical trials. Using *in vitro* experiments, it is not possible the create the full environment seen within patients and there exists no single *in vivo* animal model which completely encapsulates the pathophysiology seen within humans [9]. *In silico* models, of both mathematical and computational form, could provide a compromise: allowing us to simulate the disease in a full (synthetic) environment at a fraction of the time and cost required for *in vitro* and *in vivo* models. The ability to create a simulation model that reflects the full spectrum of pathophysiology seen in humans with TB would reduce the use of animal models and allow us to make predictions as to the efficacy of novel regimens, and thus better inform our decisions with regard to prioritisation of these new treatments. Furthermore, the development of these models allows us to explore the dynamics of TB infection and provides insight into the complex dynamics that occur and which we do not fully understand yet.

TB infection begins with the inhalation of one or more *Mycobacterium tuberculosis* (*M. tuberculosis*) bacteria, which land at the alveolar tissue of the lungs, where infection begins[10]. There, a complex battle between the host immune response and the pathogen occurs. Figure 1 shows the possible outcomes of a TB infection. In an unknown percentage of people, the immune response is sufficient and the bacterial load is low enough that the infection is cleared from the body [11]. In the rest of those infected, the bacteria proliferate and the innate immune response is insufficient to cope with the bacteria, and thus an adaptive immune response is triggered. In patients whose immune system is compromised in some manner, this adaptive immune response is also insufficient and thus active disease is formed, termed ‘primary TB’ as it originates from the initial bacterial load. This occurs in approximately 10% of those infected [12]: for the majority of patients, the adaptive immune response is strong enough to contain but not eradicate the bacteria, and the infection remains in an asymptomatic ‘latent’ form [5]. This latent form of disease represents a reservoir for bacteria [13], as the infection may re-activate if the immune system later weakens [12] and the structures containing the bacteria suffer degradation, allowing the bacteria to replicate extracellularly. This is termed ‘post-primary TB’.

**Figure 1:**
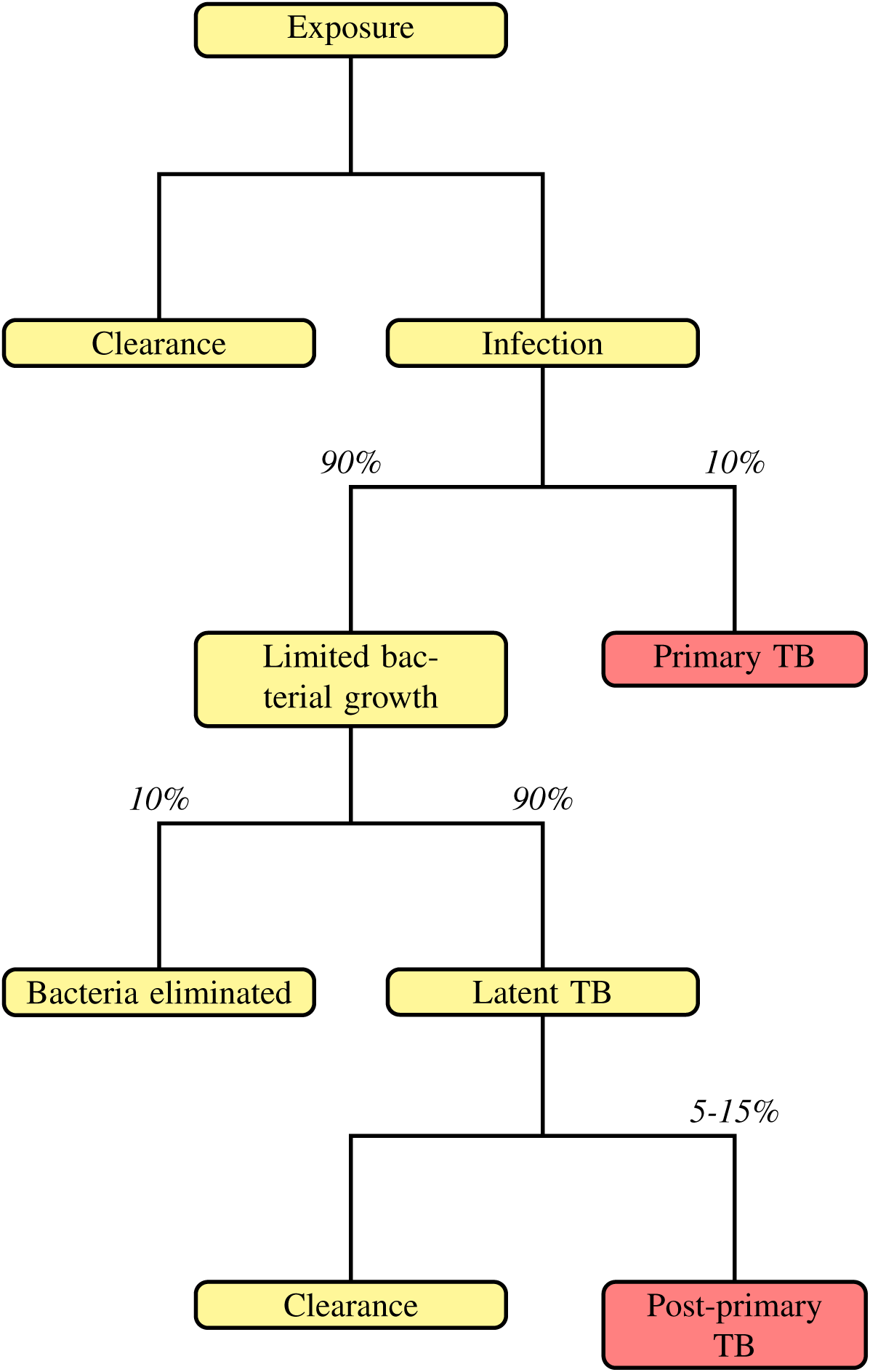
Overview of the clinical outcomes of *M. tuberculosis* exposure. Adapted from [12], values from [1, 12]. Red boxes indicate active, symptomatic disease.

Whilst both primary and post-primary stages are active, symptomatic forms of TB which can lead to patient mortality, it is important to recognise that these two forms of disease are distinct in localisation and pathology [14]. The initial infection site for TB can be anywhere in the lungs [15], but tends to occur in the lower regions [10], which are more ventilated than other lung regions [16]. Post-primary disease involves cavitation, whereby the alveolar tissue is eroded and access is permitted to the bronchial tree, and this cavitation always occurs at the apices of the lungs [15]. Cavitation does not occur during primary disease [17, 14, 18]. Thus, the bacteria that land in the lower regions must somehow disseminate to the apical regions, and the environment at the apex of the lungs must be preferential for cavitation to occur [19, 14]. It has been hypothesised that the environmental conditions within the lung contribute to these differences in localisation [15, 20, 19, 21], with factors such as the lower blood perfusion and higher oxygen tension at the apices compared to the basal regions [22, 23] believed to contribute to *M. tuberculosis* proliferation there. But exactly how each of these factors influences the progression of the disease is not well understood.

One way of incorporating environmental factors such as those described above is through the use of *in silico* models, whereby mechanistic models attempt to simulate the disease in a synthetic environment in the hopes of gaining important insight into the underlying dynamics that influence the containment or proliferation of bacteria, thus enabling us to guide treatment development towards those factors. Recently, these types of models have seen greater uptake for modelling TB (see [24] for a comprehensive review of *in silico* modelling applied to within-host TB dynamics). Early models focussed on the development of a single lesion [25], later extending to incorporate other factors such as T cell priming in lymph nodes [26], the impact of cytokines [27] and antibiotics [28]. Furthermore, models have been developed that incorporate spatial features, both within a single lesion-understanding how oxygen and chemotherapeutic concentrations within the lesion create differential regions of bacterial growth [29], and across the whole lung, investigating the dissemination of bacteria between lesions in different regions of the lung [30]. In [31], we presented the first *in silico* model of TB over the whole lung to incorporate the environmental heterogeneity present within the organ in order to understand how the differentials in factors such as blood perfusion and oxygen tension impact disease. In this paper, we build upon this model and present a new iteration of the model which includes a more granular network as its base and incorporates the full life-cycle of disease, to show how the environment within the lung impacts each stage of disease in a unique manner.

## 2. Whole lung model of TB with environmental heterogeneity

The model, termed *TBMetapopPy*, simulates the interaction between immune cells, bacteria, and the local environment both within the lung and in the associated lymphatics. It takes the form of a networked metapopulation: multiple patches exist within the model, which are linked together with edges to form a network. Patches contain sub-populations of various immune cells or bacteria, which may interact with each other and the local environment within the patch, or may translocate from one patch to another.

### 2.1. Environment

The spatial domain of the model consists of the alveolar tissue within the lung and the lymph nodes draining the lungs. The lung tissue is divided into multiple patches, each representing the total alveolar tissue present in a branch of the bronchial tree. Each patch within the lung contains environmental attributes that reflect the initial conditions within the lung at that position. These are listed in Table 1. As *V* and *Q* represent fractions of the total ventilation and perfusion, respectively, supplied to the lung that reach a patch, the sum of each of these values across all patches equals 1. The lymphatic patch contains no environmental attributes.

**Table 1:** Environmental attributes within the lung patches of *TBMetapopPy*.

| Label | Attribute | Description |
| --- | --- | --- |
| $V$ | Ventilation | The fraction of inhaled air that is passed to that area of the lung |
| $Q$ | Perfusion | The fraction of all blood sent to the lung that reaches the given patch |
| $O$ | Oxygen tension | Oxygen tension remaining in the air of the lungs after gas exchange has occurred. This is dependent on both the amount of air received ($V$) and the amount of blood received ($Q$). |
| $G$ | Drainage | The rate at which cells are able to transfer from the lung to the lymphatics system relative to the lung average. |

All patches contain populations divided into compartments, each of which represents the species and status of immune cells or bacteria, as described in Table 2. We model 4 types of bacteria, based on their location and/or replication rate. Bacteria that are present in or on the tissue surface are ‘extracellular’. *M. tuberculosis* has been shown to exhibit ‘dormancy’, associated with the accumulation of lipid bodies, whereby it reduces its replication rate but becomes more resistant to antibiotics [32, 33]. In order to incorporate this, we allow bacteria to switch between a ‘replicating’ (*B*_*ER*_) and a ‘dormant’ (*B*_*ED*_) state when extracellular. *M. tuberculosis* has evolved to suppress the destructive mechanism of immune cells by preventing phagolysosome biogenesis, and thus is able to reside within the intracellular matrix of host cells [34, 35, 36]. We define two types of intracellular bacteria - those within dendritic cells (*B*_*ID*_) and those inside macrophages (*B*_*IM*_), as both of these cells have been shown to become infected by *M. tuberculosis* [37]. We do not make the distinction between replicating and dormant here: we assume that dendritic cells are too small to permit internal bacterial replication, and that the internal environment within a macrophage is hostile and thus forces the bacteria into a slower-replicating state regardless of replication rate when ingested, as intracellular bacterial replication rates (0.1-0.44 per day) [38] have been shown to be similar to dormancy replication rates (0.25-0.5 per day) [29, 39].

**Table 2:** Population compartments within TBMetapopPy.

| Label | Compartment |
| --- | --- |
| $B_{ER}$ | Bacterium extracellular - replicating |
| $B_{ED}$ | Bacterium extracellular - dormant |
| $B_{ID}$ | Bacterium intracellular - dendritic |
| $B_{IM}$ | Bacterium intracellular - macrophage |
| $D_I$ | Immature dendritic cell |
| $D_M$ | Mature dendritic cell |
| $M_R$ | Resting macrophage |
| $M_F$ | Infected macrophage |
| $M_A$ | Activated macrophage |
| $T_N$ | Naïve T cell |
| $T_A$ | Activated T cell |
| $C$ | Caseum |

We model 3 types of immune cells: dendritic cells, macrophages and T cells. The primary role of dendritic cells in the model is antigen-presentation: the immature dendritic cells (*D*_*I*_) resident in the lung encounter and ingest bacteria, causing them to convert to a mature state (*D*_*M*_). These mature cells can then trigger an adaptive immune response by transferring to the lymphatics and activating T cells there. Macrophages play a similar role, as they can also encounter and ingest bacteria and transfer to the lymphatics, but we assume that macrophages are less mobile than dendritic cells and have a greater internal capacity - they are more likely to remain in the lung and attempt to eliminate bacteria there. Thus, macrophages have a ‘resting’ (*M*_*R*_) and ‘infected’ state (*M*_*F*_). We also include an ‘activated’ state (*M*_*A*_), whereby the macrophage’s bactericidal ability is improved during an adaptive immune response [40].

Naïve T cells (*T*_*N*_) are present within the lymphatic system and may be activated by antigen-presenting cells. Multiple varieties of activated T cells exist within the lungs with varying different functions [41, 42]. For simplicity, we do not make a distinction between the different roles and include just one type of activated T cell (*T*_*A*_) to serve as a representation for all real-world types.

### 2.2. Initial conditions

At the start of a simulation using *TBMetapopPy*, a network is constructed, using the parameters in Table 3, which models the bronchial tree as a simple space-filling tree. Firstly, a set of coordinates which define the perimeter, *P*, are provided, creating a two-dimensional shape, *S*. A point, *a*, on the perimeter is also chosen. The following algorithm is then executed to build a space-filling tree:

**Table 3:** Parameters for constructing the environment of *TBMetapopPy*. Where parameter range is Normal(*x, y*), *x* denotes mean and *y* denotes standard deviation. Where range is Uniform(*x, y*), *x* denotes minimum value and *y* denotes maximum value.

| Symbol | Description | Value / range |
| --- | --- | --- |
| $P$ | A series of $(x,y)$ coordinates composing the external perimeter of the model environment | (50, 0), (0, 0), (0, 100), (50, 100) |
| $a$ | An $(x,y)$ point, on $P$, where the branching process originates from | (50, 50) |
| $f$ | The length of the new branch as a fraction of the line dividing the perimeter in two | 2 |
| $z$ | The minimum area needed to stop the branching process | 0.05 |
| $S_V$ | The skew of ventilation values from base to apex, i.e. how many times greater the $V$ value will be for a patch at the very base of the lung compared to one at the very apex of the lung | Normal(2, 0.1) |
| $S_Q$ | The skew of perfusion values from base to apex | Normal(3, 0.1) |
| $S_G$ | The skew of drainage values from base to apex | Uniform(1, 5) |

1. A point, *b*, on *P* is chosen such the line *ab* bisects the shape *S* evenly
2. A point, *c*, is created on the line *ab*, whereby the line *ac* is a fraction, *f*, of the distance of the line *ab*.
3. Steps 1 and 2 are repeated twice, this time using *c* as the start point and perimeter series of *c* − *a* − *P*_1_ − *b* − *c* and *c* − *b* − *P*_2_ − *a* − *c* respectively, where *P*_1_ represents the points between *a* and *b* on perimeter *P*, and *P*_2_ is the points from *b* back to *a*.
4. Each iteration creates two new shapes of half the size of the parent shape. A threshold, *z*, is provided, and once the child shape sizes drop below this threshold, the iteration at that branch terminates.
5. The endpoints of the branch form the patches of the finished network.

This process (an example of which is shown in Figure 2) creates a space-filling tree: the end points of the tree may be geometrically close to one another but distant on the bronchial tree. A further patch is added to represent the lymphatic system. In order to create a network, edges are added to show translocation of members across the system. An edge is added between every alveolar patch and the lymph patch.

**Figure 2:**
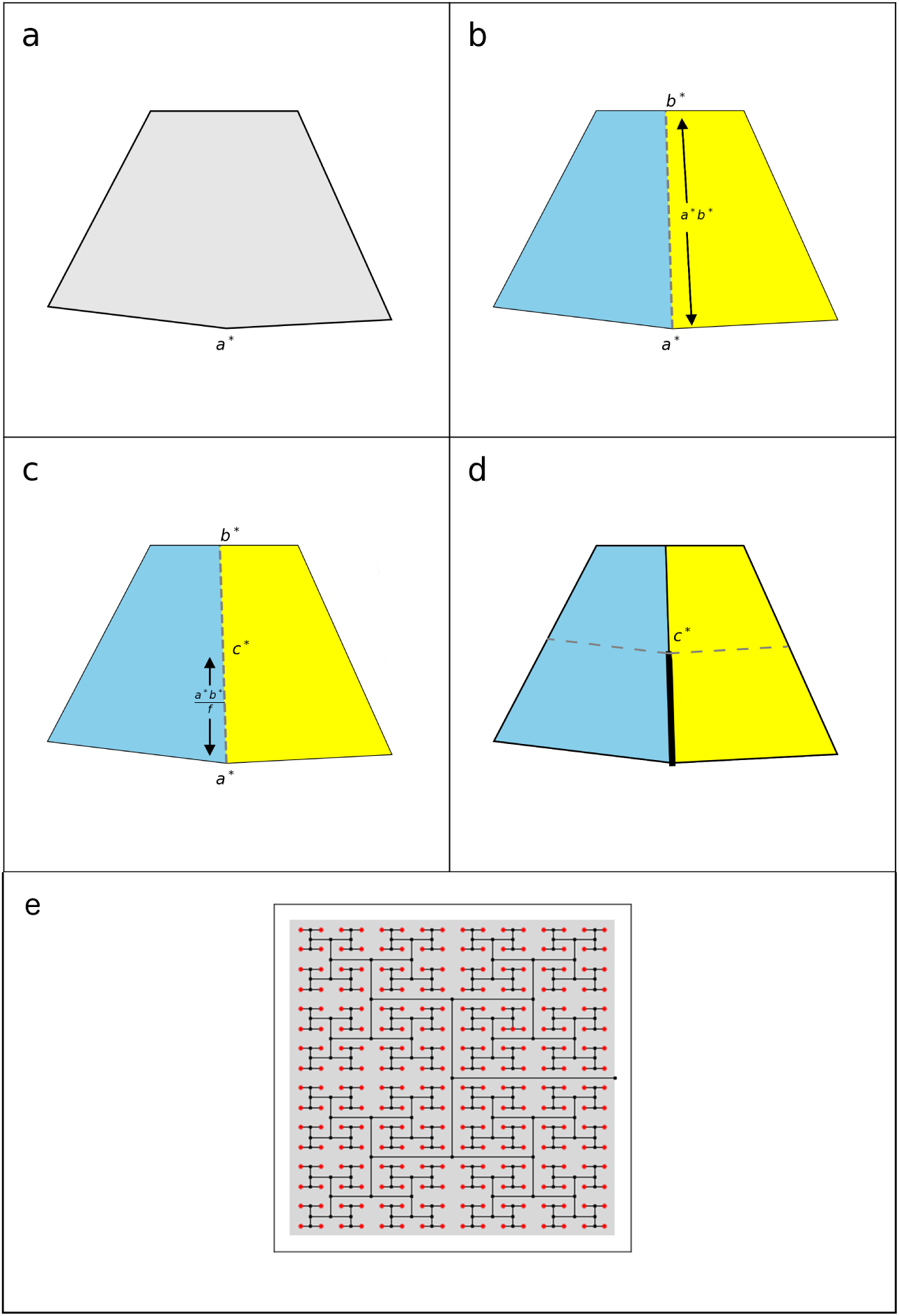
Example of the branching algorithm used to generate the environment within the second *TBMetapopPy* model. a) A shape is specified along with a branching point, *a*^∗^. b) A second point on the perimeter is chosen, *b*^∗^, such that the line *a*^∗^*b*^∗^ equally divides the shape into two equal sized shaped (coloured yellow and blue). c) A point, *c*^∗^ is chosen at a distance of *r* along the lie *a*^∗^*b*^∗^. d) A line is drawn from *a*^∗^ to *c*^∗^. The process then continues, with two new shaped (yellow and blue), and a branching point, *c*^∗^. e) The branching process applied to a rectangular shaped lung. For visual clarity, we show only the first 9 levels of the branching tree.

Once constructed, the environment is firstly seeded with values for the environmental attributes of the lung patches. Ventilation skew (*S* _*V*_) and perfusion skew (*S* _*Q*_) parameters are defined such that a patch at the very apex of the lung would have both *V* and *Q* values set to 1, whilst a patch at the very base of the lung would have a *V* value of *S* _*V*_ and *Q* value of *S* _*Q*_. Each patch, *i*, is then assigned values 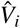 and 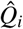 according to these scales and based on their vertical position, as shown in Equations 1 and 2, where *y*_*i*_ is the vertical position of the patch, *y*_*max*_ is the y-coordinate of the highest vertical point of the area and *y*_*min*_ is the y-coordinate of the lowest vertical point. These equations create linear scales of perfusion and ventilation across the lung, as seen in the literature [22].

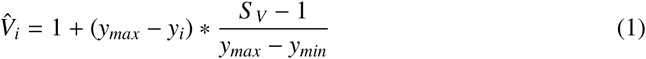

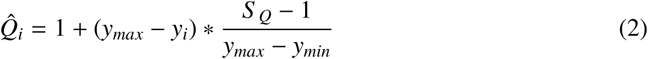

Once every patch in the lung has been assigned these values, they are normalised based on the values of each patch in the lung, as per Equations 3 and 4, which ensures the sum of all values across the lung comes to 1.

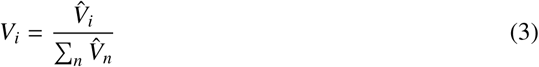

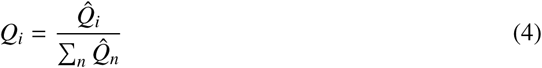

Oxygen tension, *O*_*i*_, at patch *i* is calculated automatically as per Equation 5.

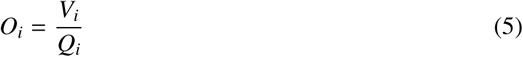

All patches (lung and lymph) are then seeded with population values. Lung patches are assumed to contain *M*_*R*_ and *D*_*I*_ in the absence of infection, with the lymph patch containing *M*_*R*_ and *T*_*N*_. We assume these populations in the lymph remain at an equilibrium level without infection calculated as per Equations 6 and 7, where *M*_*R*_ and *T*_*N*_ are the equilibrium values for the compartment *M*_*R*_ and *T*_*N*_ respectively, and other parameters are listed in Table 4.

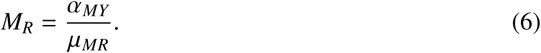

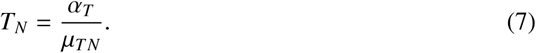

**Table 4:**
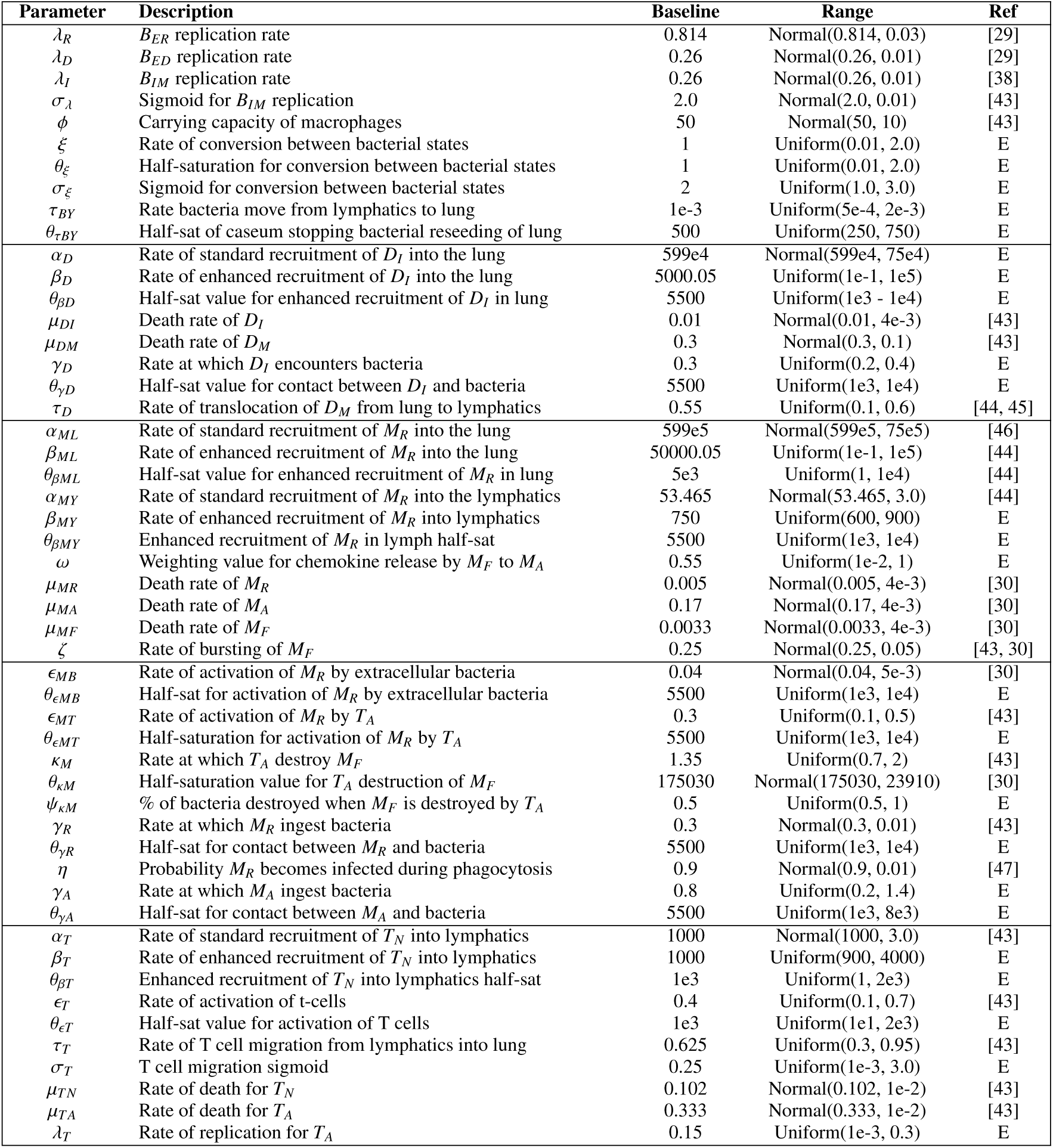
Event parameters (E = estimated, see Appendix A). All values are based on rates of event per day. Where parameter range is Normal(*x, y*), *x* denotes mean and *y* denotes standard deviation. Where range is Uniform(*x, y*), *x* denotes minimum value and *y* denotes maximum value.

Within the lung, the equilibrium value for a given compartment will be scaled by the perfusion, Q, at the patch, resulting in Equations 8 and 9.

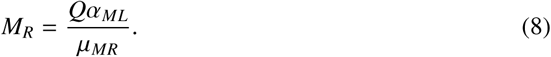

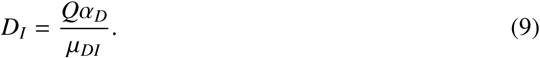

Finally, bacteria are seeded within the lung, with loads of *I*_*R*_ for *B*_*ER*_ and *I*_*D*_ for *B*_*ED*_ placed within a single patch in the lung.

### 2.3. Events

As the simulation runs, populations are able to interact with each other and the local environment, with members switching into different compartments as they change states or locations. Here, we define the dynamics that occur within the *TBMetapopPy* model. The event system of the model uses the Gillespie Algorithm to model time - each interaction is coded as a separate event and each must define how its rate of occurrence is calculated from the event parameters (listed in Table 4 and explained in Appendix A) and the current population counts of the network, and must also define the outcome of the event being performed.

#### 2.3.1. Innate immune response

*M*_*R*_ cells are initially present in both the lung and lymphatic system. These cells die naturally at a rate of *µ*_*MR*_, and are replaced through recruitment which occurs at rate *Qα*_*ML*_ in the lung and *α*_*MY*_ in the lymph. As *Q* is the perfusion value at the patch, which is itself a fraction of the perfusion to the whole lung, the parameter *α*_*ML*_ represents the total number of cells recruited to the entire lung, a portion of which will be sent to a specific patch.

*D*_*I*_ cells also die (at rate *µ*_*DI*_), and are replaced in the lung through recruitment (at rate *Qα*_*D*_). In the lymphatics, naïve T cells die (at rate *µ*_*T N*_) and are replaced through recruited cells (at rate *α*_*T*_).

The bacteria in the lung are initially extracellular and are able to replicate freely, with *B*_*ER*_ replicating at rate *λ*_*R*_*B*_*ER*_ and *B*_*ED*_ replicating at rate *λ*_*D*_*B*_*ED*_. In order to model the effects of hypoxia on bacterial replication, we include an event to switch bacteria to *B*_*ED*_ with increased likelihood in an oxygen-poor environment, with rate 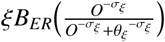, whereby *σ*_*ξ*_ > 0. We also include the reverse event, with *B*_*ED*_ converting to *B*_*ER*_ when presented with an oxygenated area, with rate equal to 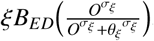.

The phagocytes within the lung may encounter the bacteria and ingest them. This occurs at rate 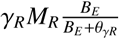 for *M*_*R*_ and rate 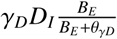 for *D*_*I*_, where *B*_*E*_ = *B*_*ER*_ + *B*_*ED*_. When performed, an extracellular bacteria is probabilistically chosen to be ingested based on current levels. We model the ability of *M. tuberculosis* to avoid destruction by assigning a probability to bacterial destruction when *M*_*R*_ encounter bacteria, with η representing the probability that a *M*_*R*_ cell will become infected and the bacterium will become intracellular, whilst (1 − *η*) represents the probability that the bacterium will be destroyed. We assume that *D*_*I*_ are incapable of bacterial destruction and so always convert to *D*_*M*_ when ingesting bacteria, and that the different replication phenotypes (*B*_*ER*_ or *B*_*ED*_) have the same chances of survival under phagocytosis.

If a bacteria survives the ingestion process, it converts to an intracellular state (*B*_*IM*_ or *B*_*ID*_ for phagocytosis by *M*_*R*_ or *D*_*I*_ respectively) and the cell converts to an infected state (*M*_*R*_ convert to *M*_*F*_, *D*_*I*_ convert to *D*_*M*_). *M*_*F*_ and *D*_*M*_ both die naturally, at rates *µ*_*MF*_ *M*_*F*_ and *µ*_*DM*_ *D*_*M*_ respectively, and this may return some of the internal bacteria back into extracellular compartment *B*_*ED*_ (we assume that the internal conditions within a macrophage are stressful to bacteria and thus they have been forced into a dormant state, which they remain in once released). For *D*_*M*_ death, we assume the single internal *B*_*ID*_ is always released, whilst for *M*_*F*_ death, we assume a percentage of the bacteria inside the macrophage 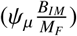 are destroyed with the remainder 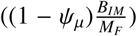 released.

Once in the internal compartment of the immune cell, bacteria can replicate, constrained by the internal capacity of the cell. We assume *D*_*I*_ cells are too small to permit replication and thus *B*_*ID*_ do not replicate. *B*_*IM*_ may replicate at rate 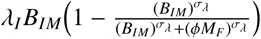. Thus the rate of replication decreases as the levels of bacteria approach the total carrying capacity of all infected macrophages (*ϕM*_*F*_). As macrophages start to fill with bacteria, they may ‘burst’, rupturing their outer wall and releasing bacteria. The rate of occurrence for this is inverse to replication, i.e. it increases as the bacterial level approaches capacity, with rate 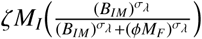. When this occurs, a fraction of total *B*_*IM*_ (calculated as 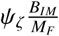) are destroyed, with the remainder, 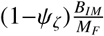, returning to the compartment *B*_*ED*_. The death of an infected macrophage releases a portion of caseum, *C*, into the extracellular compartment.

Resting macrophages may overcome their inability to destroy bacteria by activating - this converts the *M*_*R*_ member to *M*_*A*_. This process is triggered by the presence of bacteria, at rate rate 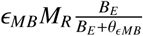. Activated macrophages can ingest bacteria in the same manner as *M*_*R*_, with rate 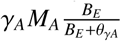, but *M*_*A*_ cannot become infected and thus the bacteria ingested are always destroyed, simulating the increased bactericidal effects of macrophage activation. *M*_*A*_ cells die at rate *µ*_*MA*_ *M*_*A*_.

Infection is not just restricted to the initial location and the lymphatics. We assume that there exists a period of time after bacteria become present in the lymphatics and before the establishment of a structured lesion there, and that during this time period bacteria are able to freely move from the lymph nodes into the blood and thus be transferred back into the lung to seed new lesions. In modelling terms, we assume that the presence of caseum indicates a stable lesion, and thus the rate of translocation of bacteria from the lymphatics into the lung is 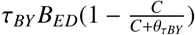.

#### 2.3.2. Adaptive response

In response to infection, the body increases the supply of immune cells to the lung to assist with containment. This is modelled by separate events for each resident immune cell type at each location. In the lung, enhanced macrophage recruitment occurs at rate 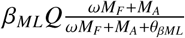. We assume that *M*_*F*_ and *M*_*A*_ cells create the necessary chemokines to trigger enhanced recruitment, and include *ω* as a weighting term to allow for these cells to produce differing levels of chemokine. Similarly, dendritic cell recruitment is enhanced at rate 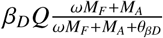. In the lymphatics, macrophage recruitment is enhanced at rate 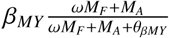. Enhanced T cell recruitment is dependent on dendritic cells and is increased at rate 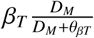.

Dendritic cells transfer to the lymphatics during infection in order to present antigens and trigger an adaptive immune response [48, 49]. We model this transfer to the lymphatics with an event that allows *D*_*M*_ to translocate from the lung to the lymphatics at rate *τ*_*D*_*GD*_*M*_, thereby creating different rates of clearance at different regions, based on *G*. This transfer, whilst necessary for establishing an adaptive immune response through the presentation of antigen, also carries the risk of spreading infection, as some immune cells may harbour internalised bacteria and these will also transferred to the lymphatics. The initial activation and proliferation of *M. tuberculosis*-specific T cells has been shown to be determined by the number of bacteria present in the lymph nodes [50], suggesting transfer of bacteria to the lymph nodes may be important for establishing an adaptive immune response. We assume that our model simulates a severe form of infection whereby bacteria are disseminated to the lungs in this manner, and thus transfer of a *D*_*M*_ member also transfers a single *B*_*ID*_.

Antigen-presenting cells (both *D*_*M*_ and *M*_*F*_) in the lymphatics can cause activation of T cells from a naïve to an activated state. This occurs at rate 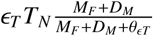, and results in *T*_*N*_ changing to *T*_*A*_. Once activated, these T cells migrate to the sources of infection to enhance the immune response and contain the bacteria. We allow the body to direct T cells to where they are needed by basing the rate of translocation of these cells from the lymphatics to the lung on the number of *D*_*M*_ - the presence of these cells in the lymphatics gives an indication of the level of infection within the lung as they only originate within the lung. Thus, increased numbers of *D*_*M*_ in the lymphatics indicates that more T cells need to migrate to contain infection in the lung, rather than remaining in the lymph to fight infection there. The rate of translocation is 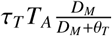.

Activated T cells perform two primary functions in the model: the first is to cause activation of macrophages, converting *M*_*R*_ into *M*_*A*_, occurring at rate 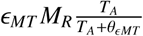. The second function of *T*_*A*_ is to causes apoptosis - the controlled destruction of infected cells. This is modelled by allowing *T*_*A*_ members to destroy *M*_*F*_ members at rate 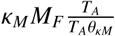. Doing so also destroys some internal bacteria, calculated as 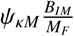, with 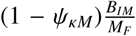 converting from *B*_*IM*_ to *B*_*ED*_. *T*_*A*_ members naturally die at rate *µ*_*T A*_*T*_*A*_.

#### 2.3.3. Weakening of the immune system

The events described previously constitute the primary and latency stages of infection: an initial infection occurs within the lungs and the bacteria levels are brought under control by an adaptive immune response. In order to model a post-primary infection (more specifically, a reactivation scenario whereby the immune system containing the bacteria during the latency phase is compromised and thus allows for bacterial levels to rise again), we include a series of events that gradually reduce the rate of T cell recruitment (i.e. *α*_*T*_ is decreased). These follow Equation 10, where

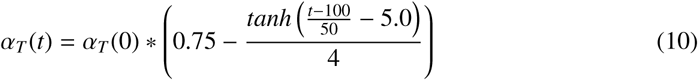

Unlike the previous events, these events (occurring at t=0,1,2,…,600) are not stochastic and instead are coded to occur at their respective time-points. For our experiments, we chose this function as it allows for a 50% reduction in T cell recruitment after approximately 250 days, a time-point where it was determined empirically that bacteria numbers in each lesion oscillate but never the levels seen in the initial lesion during primary infection, as seen in Figure 3B. Figure S1 shows Equation 10 applied to baseline value of 1000 for *α*_*T*_ (0). These events are not intended to mimic a specific disease (such as human immunodeficiency virus (HIV)), but represent an abstract weakening of the immune system, which might occur through a variety of external triggers.

**Figure 3:**
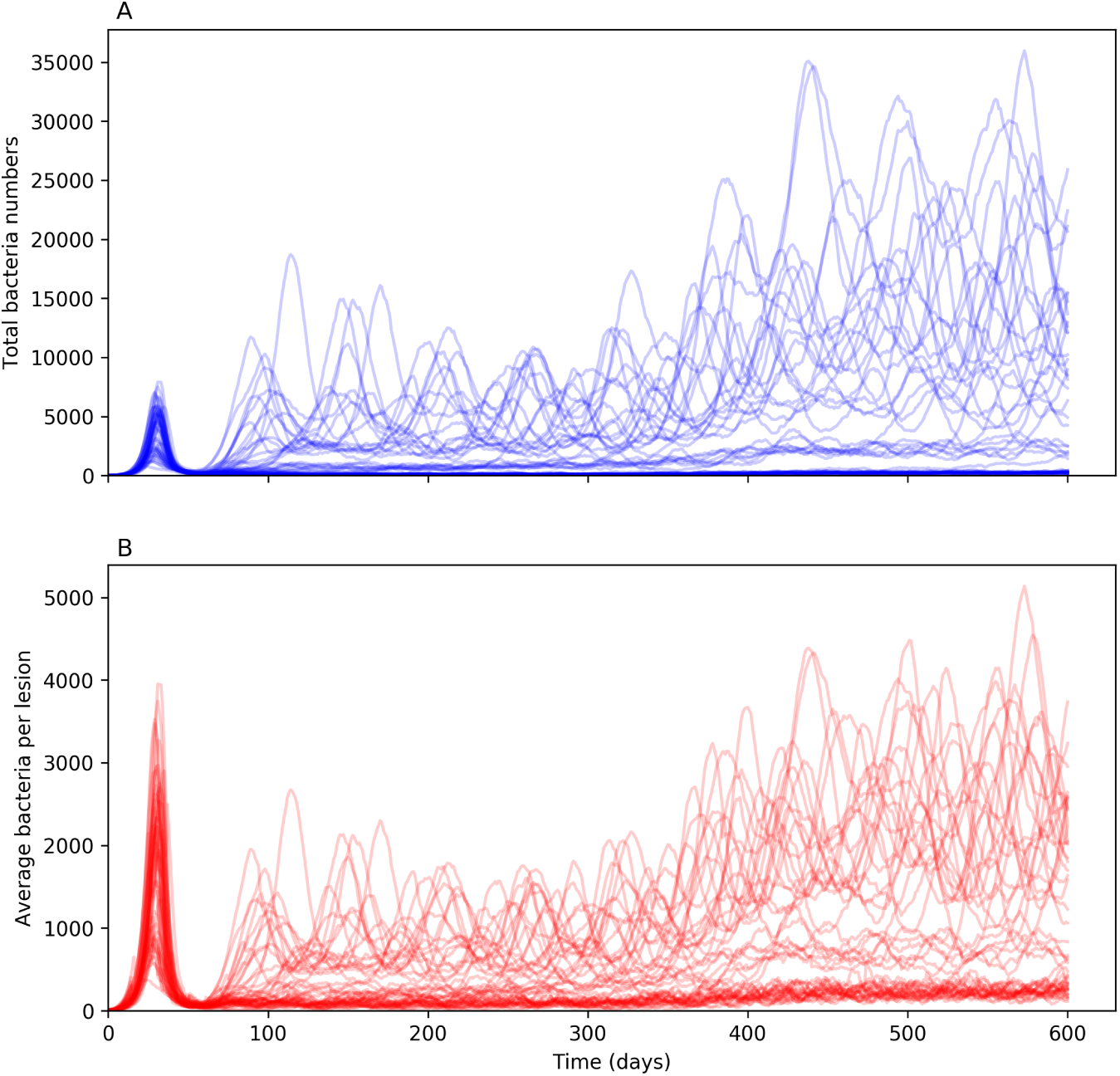
A) Total numbers of CFU within the entire system over time for each simulation of the *TBMetapopPy* system B) Average CFU count per lesion over time within each simulation. Both plots show counts for each of 50 repetitions.

## 3. Results

We began by running simulations using the model with parameters set to the baseline values as defined in Tables 3 and 4. As the model is stochastic, multiple repetitions were required - 50 repetitions were run in total.

### 3.1. Primary infection to latent infection - total bacteria and bacteria per lesion

We first examined the total bacterial load of the system over time. Figure 3A shows the results of the total bacterial load for each of the 50 repetitions over time. In the initial phase, a primary infection begins and reaches its peak between 25 and 45 days in all simulations. This is brought under control by the introduction of the adaptive immune response with low bacterial levels in all simulations by about day 55. This correlates with evidence that the adaptive immune response for TB occurs within humans within 5-6 weeks [45].

Whilst all simulations are relatively consistent to the point where the infection has been brought under control, with only minor variations in the height and time of the peak of initial infection, the results vary much greater in the response after this point. Whilst many simulations tend to keep bacteria numbers at a low level, for those that do not, the total number present has a large amount of variation. However, it is important to understand the size of individual lesions: a large bacterial load in total may represent a single lesion with a high bacterial load or multiple lesions with low loads. It is important to make the distinction between these two scenarios: multiple small lesions can be interpreted as being stable: the immune system is able to keep these lesions at a low level and stop bacterial growth; whereas a large lesion can be interpreted as a failure of the immune system to control bacterial growth at that location: this may lead to tissue damage, and then cavitation. Therefore, it is preferable for the host to keep the number of bacteria in a single lesion small, and it is important for us to understand the number of CFUs in each lesion. From here onwards, we refer to the number of bacteria at a lesion as lesion CFU.

We investigated the notion of bacteria distribution by tracking the average lesion CFU for each simulation, with the results shown in Figure 3B. Here, we see that the average lesion CFU after day 60 is much lower: indicating that while there are a high number of bacteria, they are being spread across multiple lesions resulting in a small average lesion CFU. At the time of the T cell recruitment drop (t=250 days), the average lesion CFU increases again, in many cases reaching levels similar to those seen at primary disease. We determine these to be a post-primary form of disease. We also note that not all replications reach this level: in some cases, where the average CFU count was low during latency, the count after the T cell recruitment drop increases but does not reach these high levels.

### 3.2. Spatial distribution of average lesion CFU and number of lesions

We then explored which spatial areas of the lung were being affected by the disease over the course of the infection, by looking at the average lesion CFU for each of 15 evenly-sized horizontal divisions of alveolar tissue (i.e. we group patches together that fall within the same given position boundaries based on their height within the lung). Figure 4A shows the correlation of average lesion size to vertical position in the lung during the various stages of infection. The initial lesion (in the very lowest region) grows and peaks at around 30 days. By day 50, the lesion has stabilised but bacteria have spread to other regions of the lung. At this point, these lesions are all at a very small average CFU count, despite the differences in their life-cycles: the large, older lesion at the base has almost completely healed, whilst the lesions at other regions in the lung have just begun and thus have lower CFU.

**Figure 4:**
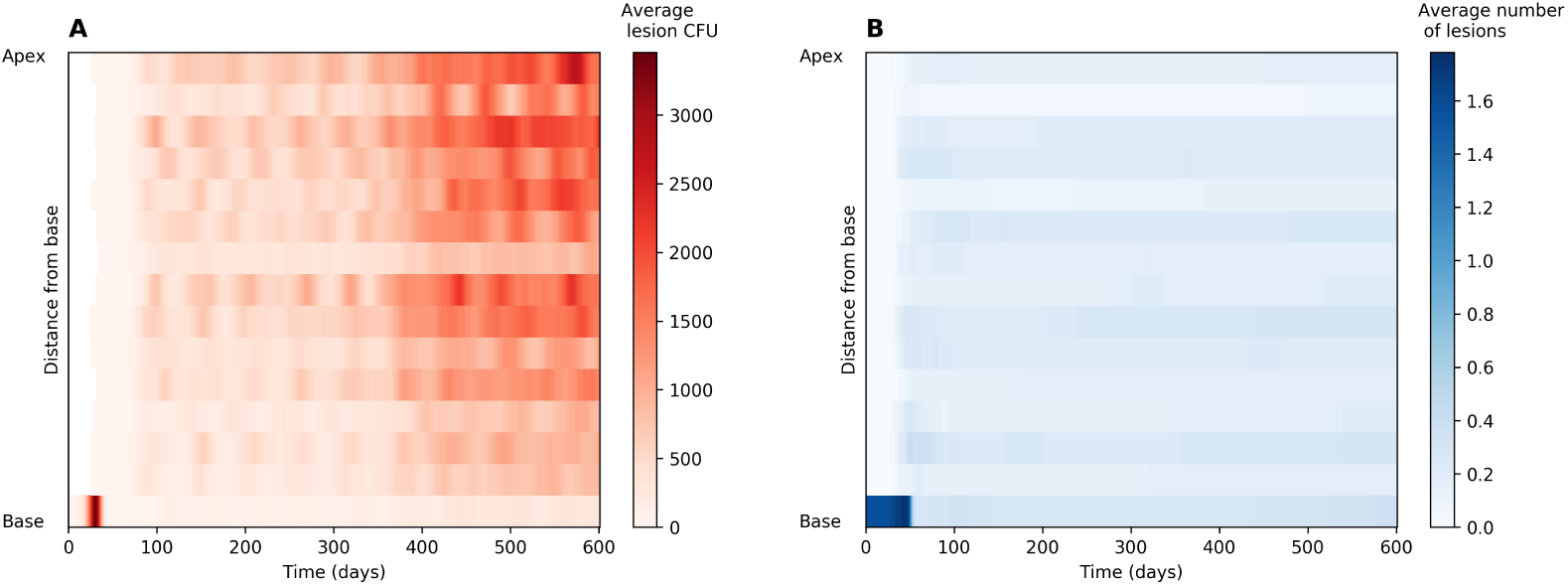
A) Vertical distribution of CFU count per lesion within *TBMetapopPy* simulations over the life-cycle of the infection. Colour shows the average CFU count at each of 15 evenly sized horizontal slices of lung taken from 50 simulations, and the progression of these averages over time. B) Vertical distribution of the average number of non-sterilised lesion per simulation within *TBMetapopPy* over the same 50 simulations.

Latency is visible in days 75 to 250. We witnessed that whilst lesion CFU count starts homogeneous across the lung, by day 100 the numbers of bacteria at lesions in the apical region start to expand at a greater rate than those towards the base. This is in stark contrast to the lesions during the primary stage, which are typically completely homogeneous in terms of bacterial load. The CFU count of these lesions during latency fluctuates in a sinusoidal manner (see Figure 3A), but these average CFU counts never reach the size seen during the primary infection.

In Figure 4A we see how the spatial distribution of average lesion CFU within the lung changes post T cell recruitment drop (see Supplementary Materials A for a breakdown of lesion sizes by each simulation). After the start of the T cell recruitment drop at 250 days, the average CFU count increases, with the largest, most damaging lesions appearing towards the apex of the lung. The lesions in the lower regions also increase in CFU but not to the levels seen towards the apex.

Furthermore, we investigated the number of non-sterilised lesions (lesions that have bacteria present) at different areas of the lung (Figure 4B). We note that at the base of the lung, the average number of lesions starts at 1 and subsequently drops, implying that the initial lesion is often sterilised. Throughout the rest of the lung, the number of lesions was relatively even and not affected by the recruitment drop - implying the weakened immune system results in existing lesions increasing in bacterial load rather than the formation of new lesions in the model.

Figure 5 shows the bacterial counts of lesions within a single simulation. Lesions are labelled according to their vertical position in the lung, either apical or basal. We note that during latency, apical lesions maintain higher bacterial loads than those towards the base. Following T cell recruitment drop, all lesions oscillate in bacterial numbers, with apical lesions experiencing greater amplitude of oscillations. Within the lymph patch, bacteria levels remain fairly constant, even after the reduction in T cell recruitment.

**Figure 5:**
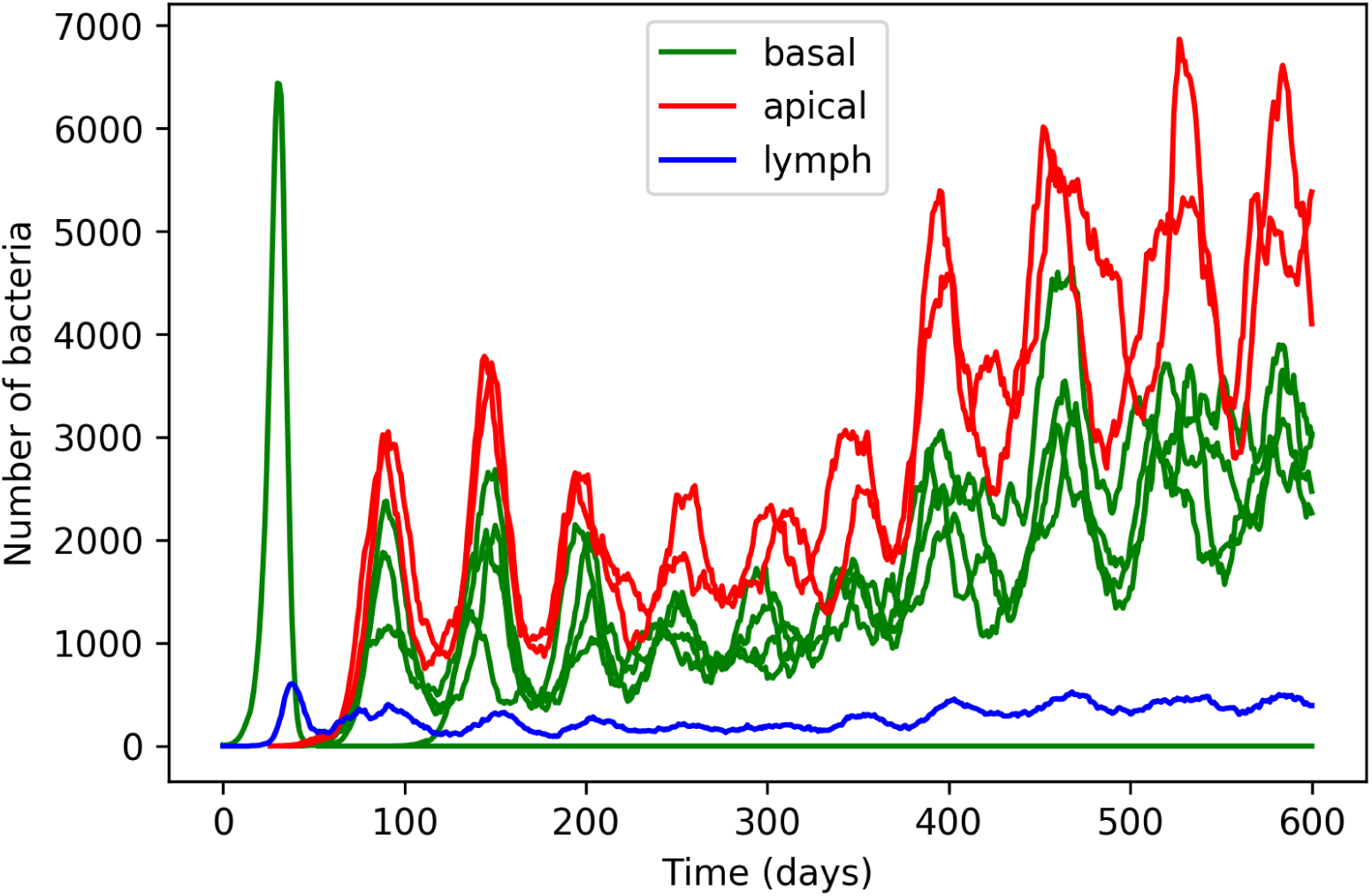
The bacterial counts of individual lesions within a single run. Lesions are labelled either apical when above the vertical centre of the lung, basal when below the vertical centre, or lymph for the lymphatic patch.

### 3.3. Lesion composition

Figure 6 shows the composition of populations of lesions in different regions at different points of time in a single simulation. At t=240 days (i.e. during latency), the bacterial composition of both lesions is predominantly intracellular, and within macrophages, with the apical lesion having a higher bacterial load. After the T cell recruitment drop, both lesions increase bacterial numbers. However, we note that the bacteria at the apex now contain a small but significant sub-population of extracellular bacteria. For immune cells, we note that the basal lesion contains greater bacterial numbers in total (due to its increased perfusion compared to the apical location). At the apical region, there is a greater population of infected macrophages, and this difference increases after the recruitment drop at t=400 days. In both cases, the total number of immune cells in the lesions decreases.

**Figure 6:**
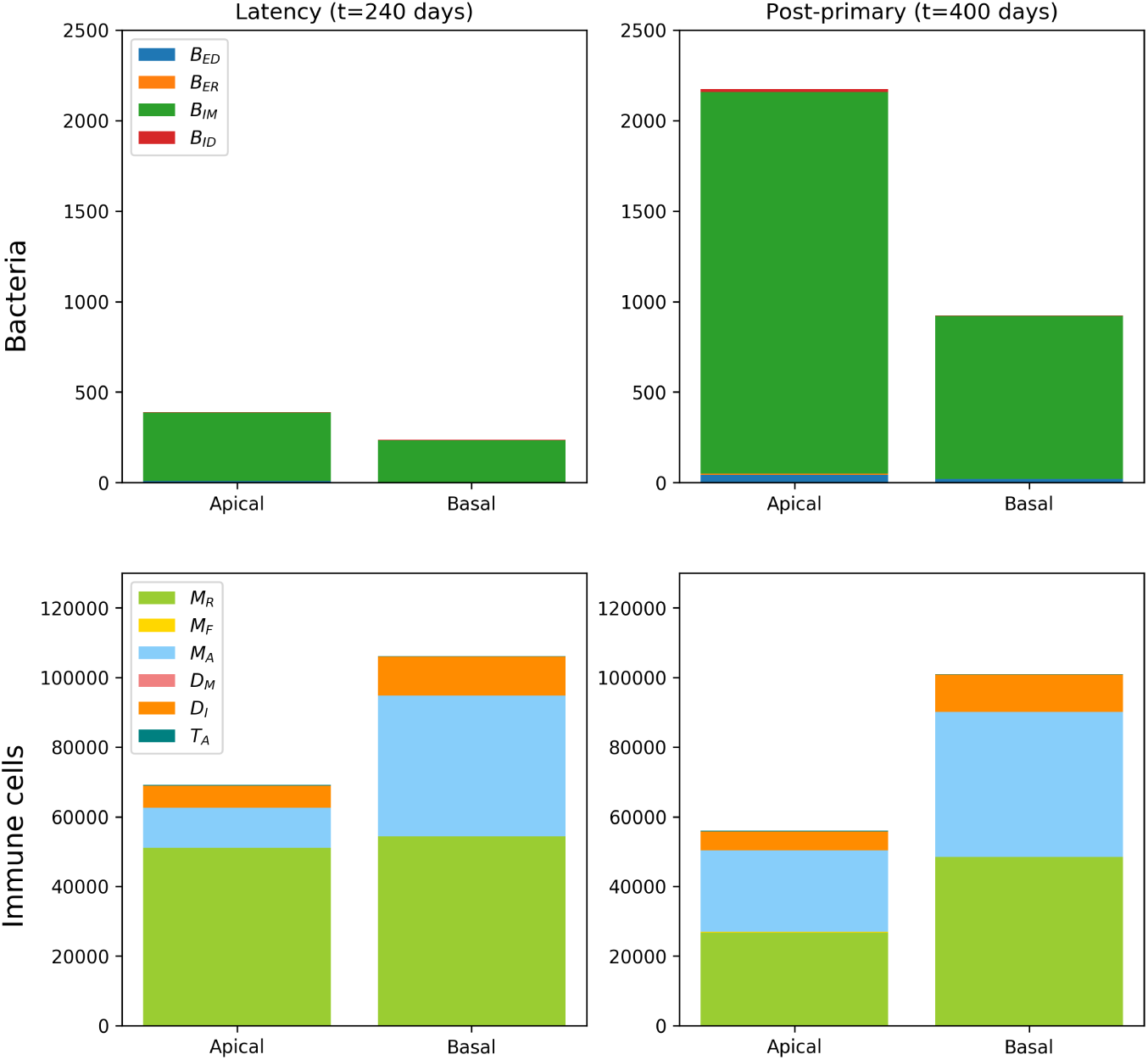
The composition of lesion populations during latency (t=240 days) and during post-primary disease (t=400 days). Two lesions from a single simulation were chosen - one towards the apex (“apical”) and one closer to the base (“basal”). Populations are separated into bacteria (first row) and immune cells (second row).

## 4. Sensitivity Analysis

Having established that the environmental heterogeneity present within the lung environment can plausibly contribute to the differences in localisation seen during different stages of infection, we then proceeded to run uncertainty and sensitivity analysis, exploring the scale of uncertainty present within the results of our model and apportioning this uncertainty to the uncertainty within our input parameters. The reasons for doing this are two-fold. Firstly, our model, like any model constructed to simulate real-world biology, is built upon real-world data which is often incomplete [51], possibly due to a lack of available *in vivo* and *in vitro* models. As such, many of the parameters used in the model have been estimated and therefore should be tested for various values to determine how the uncertainty in these parameters affects the model output. It is important to understand exactly how uncertainty in the model input propagates to its outputs, as this can highlight which uncertain parameters should be investigated further (perhaps in a laboratory or clinical trial setting) in order to reduce the parameter uncertainty and thus improve our confidence in the model’s outputs. Secondly, by varying the values of input parameters and tracking the variance of the model’s outputs, we are afforded insight into which of the model parameters is most influential on the given output, and thus can deduce which individual events within the system most influence disease outcomes. Future treatments that can target these important functions would be more likely to be successful.

Our model contains a large number of parameters based on biological processes which are uncertain, and this introduces a large amount of uncertainty into our results. As these parameters drive various events within the model that rely on the same elements (such as the bacterial and immune cell compartments), it is reasonable to assume that there will be interactions between parameters. Furthermore, the complex host-pathogen interactions that occur during TB disease mean that we can expect non-linearities to be present in our model as well, as shown in our results in Figure 4: the reduced perfusion towards the apex improves the chances of bacterial growth in that region, but the average lesion CFU does not rise towards the apex in a linear fashion. These factors of interactions and non-linearities mean that in order to perform a valid sensitivity analysis, a global approach must be used, i.e. all parameters must be varied at the same time [51], and the whole *n*-dimensional (where *n* is the number of parameters) parameter space must be explored.

In order to do this, we chose to follow the methodology laid out by Marino et al in performing global uncertainty and sensitivity analysis of systems biology models [52]. For our parameter value ranges, where parameters were derived from values from the literature, we assigned a range based on a normal distribution using the literature value as the mean and an appropriate standard deviation value. For parameters which have been estimated, a uniform distribution was chosen with a large range to reflect the additional uncertainty.

There exists a large number of possible outputs from the model. For the purposes of our analysis, we examined seven outputs, as listed in Table 5. These values vary over time, as the system moves between different stages of infection. Therefore, we track the sensitivity values of each parameter and output combination over a time period, chosen as being day 1 of infection to day 550. We track three variables, Ω_*N*_, Ω_*B*_ and Ω_*V*_, measuring total bacteria levels, lesion counts and average lesion CFU across the whole lung, and a further six variables looking specifically at the apex of the lung (which we define as being above the centre of the lung for this analysis).

**Table 5:** Outputs from *TBMetapopPy* used for sensitivity analysis.

| Output | Description |
| --- | --- |
| $\Omega_B$ | The total number of bacteria (of all states) present within the system |
| $\Omega_N$ | The total number of non-sterilised lesions present within the system (i.e. patches where bacteria count > 0) |
| $\Omega_V$ | The average lesion CFU of all non-sterilised lesions within the system (i.e. the average number of bacteria per patch, where number of bacteria > 0) |
| $\Omega_{AB}$ | The total number of bacteria at apical patches (i.e. patches above the centre of the lung) |
| $\Omega_{AE}$ | The number of extracellular bacteria in apical patches |
| $\Omega_{AL}$ | The number of lesions in apical patches |
| $\Omega_{AV}$ | The average lesion CFU of apical patches |

These track total bacteria numbers (Ω_*AB*_), as well as extracellular bacteria (Ω_*AE*_) and replicating extracellular bacteria (Ω_*AR*_). We also measure number of lesions in the apex (Ω_*AL*_), average lesion CFU in the apex (Ω_*AV*_) and number of activated T cells there (Ω_*AT*_).

### 4.1. Uncertainty analysis

We first plotted the results of our outputs in order to perform our Uncertainty analysis and understand how uncertain each output was. We chose to use 50 stratifications of the input parameter ranges, and used the average results of 20 repetitions in order to reduce the contribution of aleatory uncertainty. These are shown in Figure 7, which shows the plots of each of the 50 aggregations of 20 repetitions for each parameter sample generated by a Latin Hypercube Sampling method.

**Figure 7:**
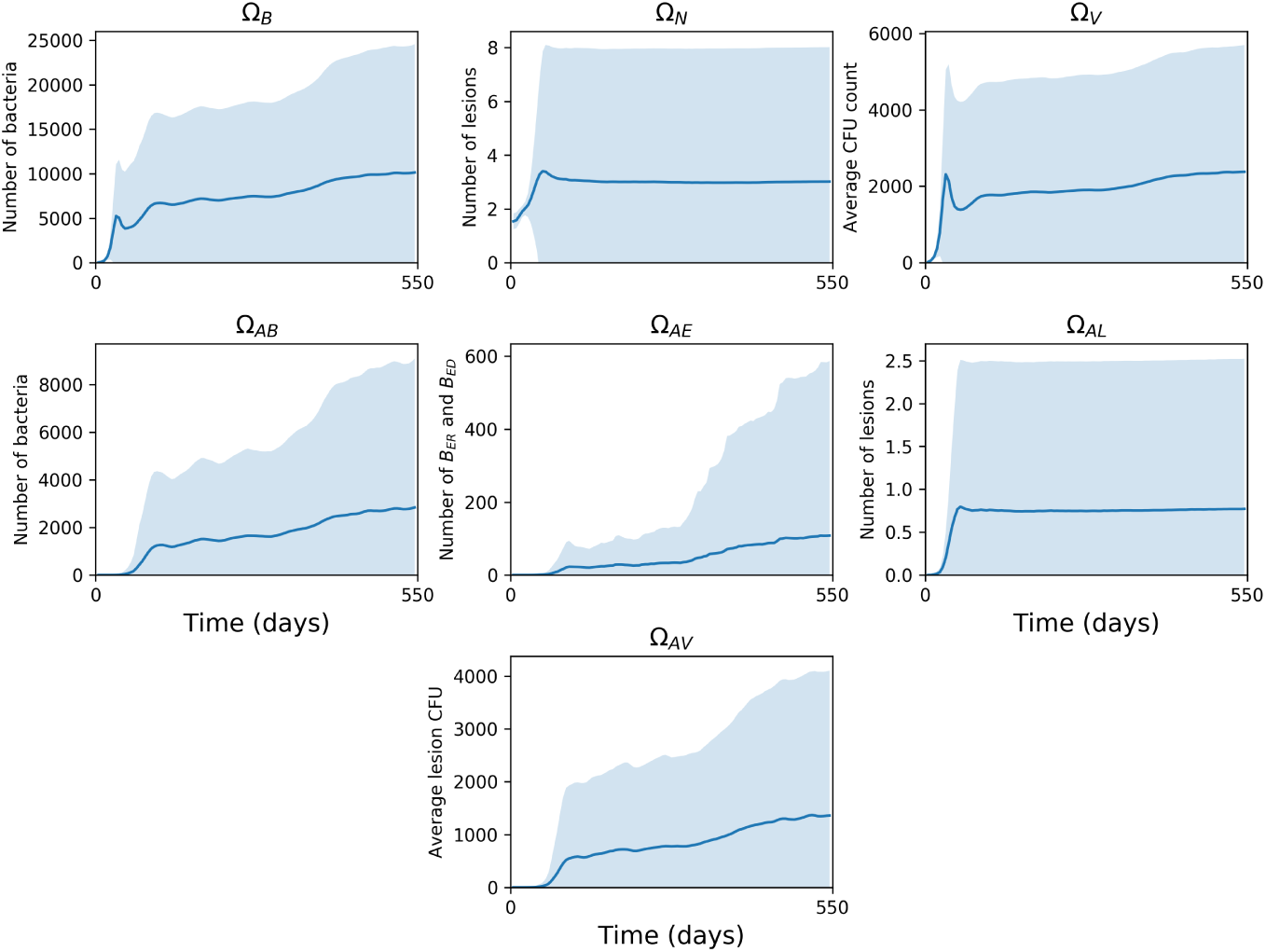
Uncertainty analysis plots of each output (listed in Table 5) over time. Each plot shows the mean (solid line) and standard deviation (shading) of the output values from each of the 50 parameter sample averages (each sample is an average of 20 repetitions)

From these plots, we note that there is a high degree of uncertainty in our output variables, particularly in the later stages of infection (as seen by the large errors present for all three outputs). For the variables tracking the whole lung, we note that the total number of bacteria and the average lesion CFU (and their uncertainties) are affected by the immune cell recruitment drop, with their numbers increasing within the post-primary phase. The number of lesions does not seem to be impacted by the drop.

Within the apex, we note that for bacterial numbers, particularly for extracellular and replicating extracellular bacteria, there is a marked increase after the T cell recruitment drop, and that the uncertainty within these parameters also greatly increases (this is especially the case for replicating extracellular bacteria). Similarly to the whole lung, the number of lesions at the apex remains constant throughout (slightly below 1),with the uncertainty ranging up to approximately 2.5 and this also does not change throughout the course of infection. The average lesion CFU increases after the recruitment drop (similarly to previously mentioned outputs).

### 4.2. PRCC

Having established the uncertainty within our model outcomes, we then generated Partial Rank Correlation Coefficient (PRCC) values for each parameter and output combination over the time-scale of the simulations. The full set of plots are presented in the Supplementary Materials Figures S3-S7, and in this section we overview some of these results.

#### 4.2.1. The influence of environmental parameters

The three environmental parameters, *S* _*V*_, *S* _*Q*_ and *S*_*G*_, all exhibit differing sensitivities upon the outputs (see Figure 8). The value of each of these variables impacts the heterogeneity in the lung, with larger values creating a greater differential between the apex and the base for the given environmental factor; for example, a greater *S* _*V*_ value results in more ventilation being directed towards the base of the lung and less towards the apex, and similarly for *S* _*Q*_ and *S*_*G*_.

**Figure 8:**
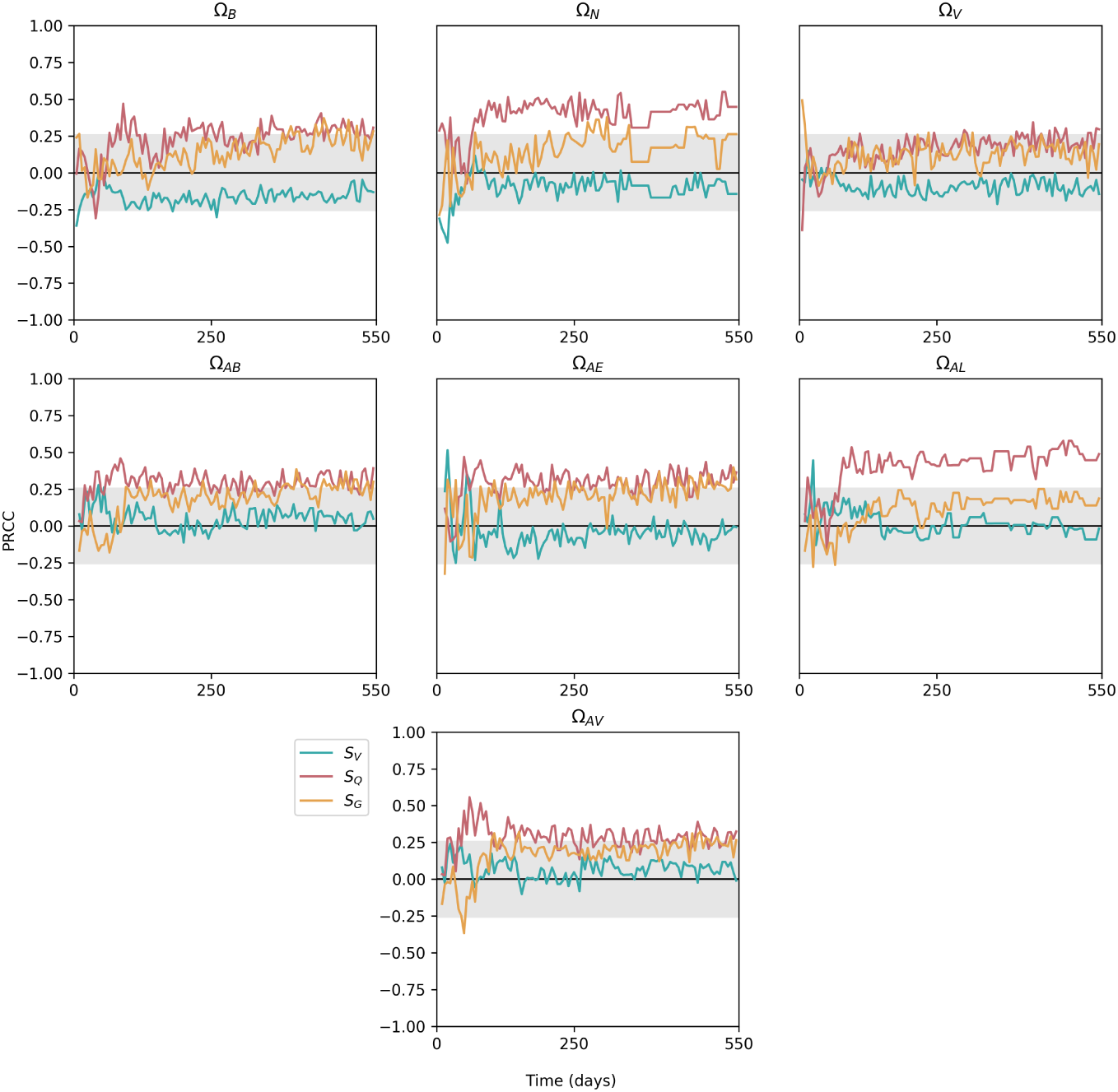
Sensitivity plots for environmental parameters within *TBMetapopPy*, plotting the PRCC value of each parameter against one of the model outputs over time. Grey shaded area shows non-significance (p<0.01).

*S* _*V*_ only has significance on the outputs during the primary stage of infection, with a negative effect on Ω_*B*_ and Ω_*N*_ at the very start of infection and a positive effect on Ω_*AE*_ and Ω_*AL*_ shortly after the start. Throughout the remainder of infection, the differential in ventilation has little effect on our outputs. However, he differential in perfusion, *S* _*Q*_, has a more pronounced effect. This variable shows positive significance for most of output variables across the whole time-frame of infection, suggesting that the greater the differential in blood supply between the apex and the base, the worse the outcome of infection. This is particularly pronounced for Ω_*AL*_ - a higher differential in perfusion leads to a larger number of non-sterilised lesions being present at the apices in later stages on TB. *S*_*G*_ follows a similar pattern to *S* _*Q*_ but often with less significance. We note that Ω_*V*_ has a relatively high sensitivity to *S*_*G*_ at the initial stage of infection, suggesting better drainage at the lower region of the lung reduces the size of the initial lesion.

### 4.2.2. The importance of T cells

Figures 9 and 10 show the sensitivities of parameters related to T cells that exhibit significance at some point in the time-frame, against the outputs of the model. We note that the model is highly sensitive to many of these parameters, indicating that T cells play a very important role in the course of infection (we do not measure sensitivity to *α*_*T*_ since this parameter does not remain constant throughout infection).

**Figure 9:**
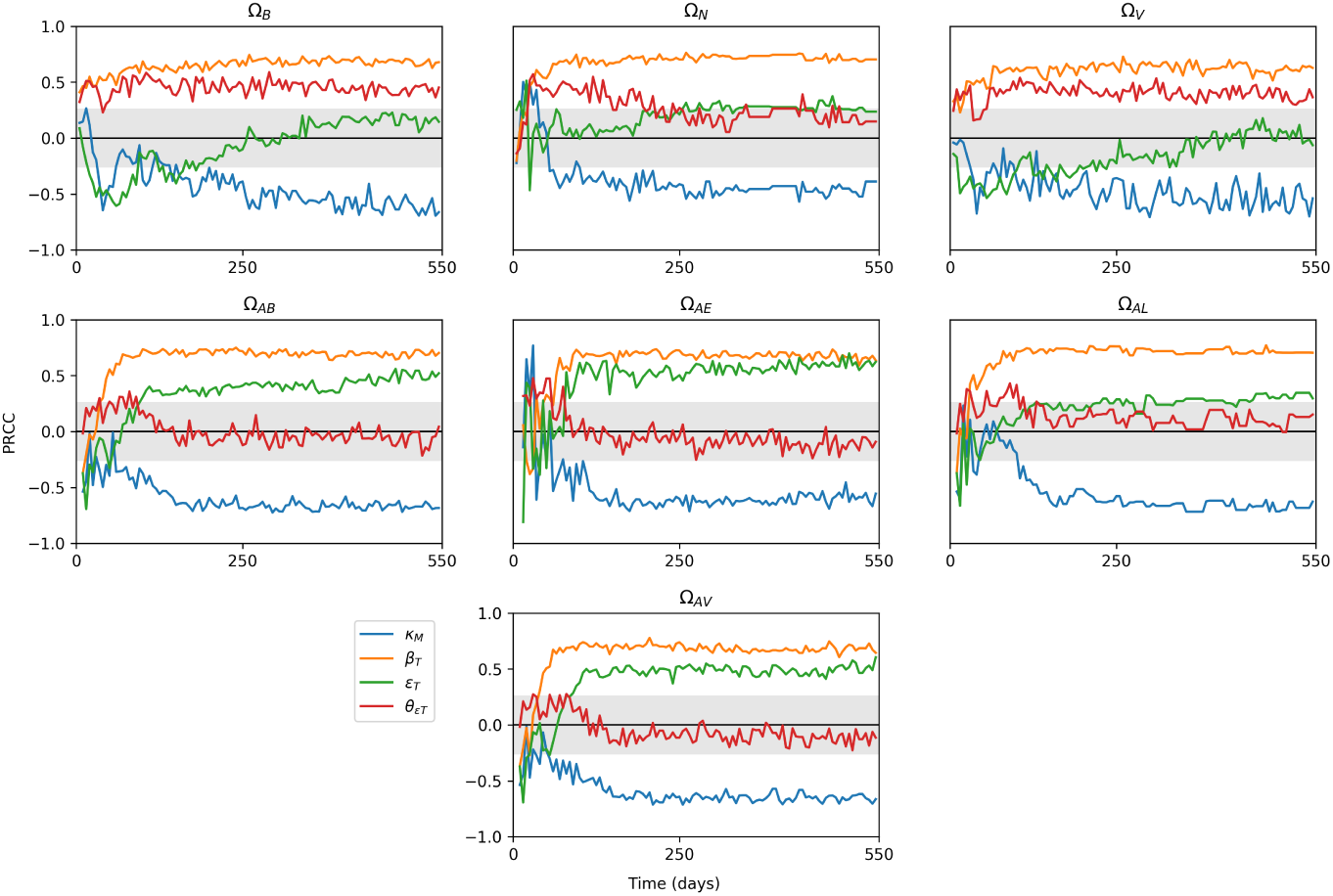
Sensitivity plots for T cell related parameters within *TBMetapopPy*, plotting the PRCC value of each parameter against one of the model outputs over time. Grey shaded area shows non-significance (p<0.01). Only parameters with sustained significant PRCC values against one output are displayed.

**Figure 10:**
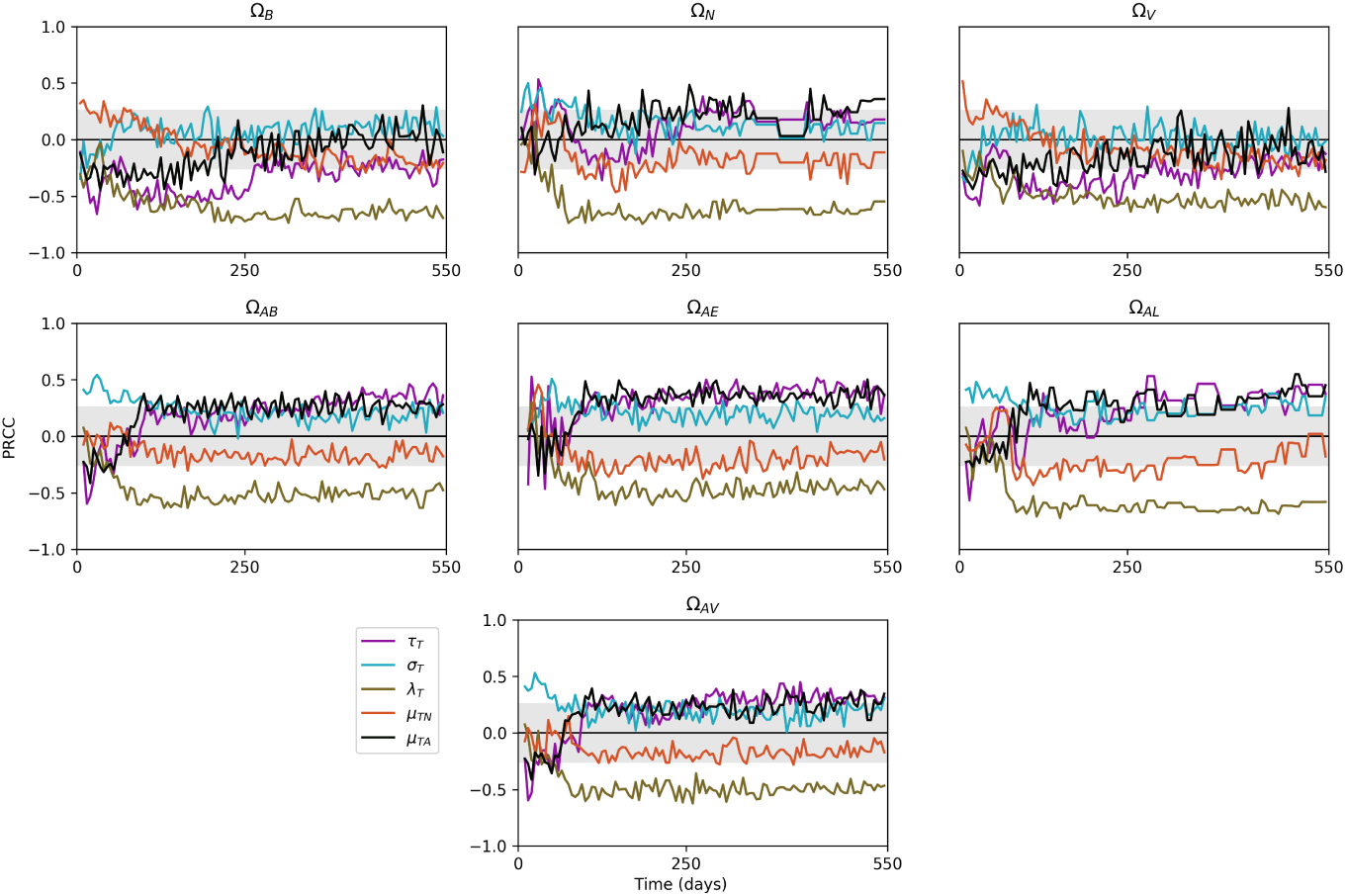
Further sensitivity plots for T cell related parameters within *TBMetapopPy*.

From Figure 9, the rate of destruction of macrophages by T cells *κ*_*M*_ has a significant impact on all outputs for the majority of infection. In the initial stages, this impact is positive for Ω_*N*_, suggesting better destruction of macrophages may be the cause of initial seedings of secondary lesions, as it releases bacteria from the internal compartment and allows them to disseminate (this is also seen in the positive sensitivity of Ω_*AE*_ to *κ*_*M*_). However, as infection progresses, these positive sensitivities become negative, suggesting that whilst destruction of infected macrophages during the early stages may help spread infection, in post-primary disease it is an effective containment measure. Interestingly, enhanced recruitment of T cells due to infection (*β*_*T*_) and the rate of T cell activation (*ϵ*_*T*_) both positively impact bacteria numbers throughout the lung and at the apex.

From Figure 10, the replication of activated T cells has a strong negative impact on all variables for most of the life-cycle, with greater values after the primary infection has been established. Conversely, the death rate of activated T cells (*µ*_*T A*_) typically has positive impact, implying faster death of T cells weakens the fight against the bacteria, although this impact is negative during the primary stage. These findings suggest that while a strong T cell response is necessary during post-primary infection, too strong a response in the primary stage could have a negative impact, possibly by releasing bacteria into the extracellular compartment and allowing them to disseminate before a solid lesion in the lymphatics can be formed.

#### 4.2.3. Bacterial replication

Our PRCC results show contrasting findings for the replication of intracellular and extracellular bacteria on the outputs, as shown in Figure 11. Of the three bacterial phenotypes which replicate in the model, it is *B*_*ED*_ whose replication has the greatest significance on the model outputs. During primary infection, this is a significant negative impact, implying replication of dormant bacteria may be driving an immune response that quickly contains infection. Thus, it is preferential for bacteria to build slowly, and thus avoid triggering too strong a response from the host before dissemination of bacteria can occur. This replication then switches to positive during later stages of infection. At the apex, extracellular replication of replicating bacteria (*λ*_*R*_) has a negative impact on extracellular bacteria numbers. This may be a cause of oscillations seen in bacteria numbers: replication of extracellular bacteria results in an increase of T cell migration to the area, which ultimately brings bacteria numbers down. This continues in a cycle with bacteria and T cell numbers both rising and falling.

**Figure 11:**
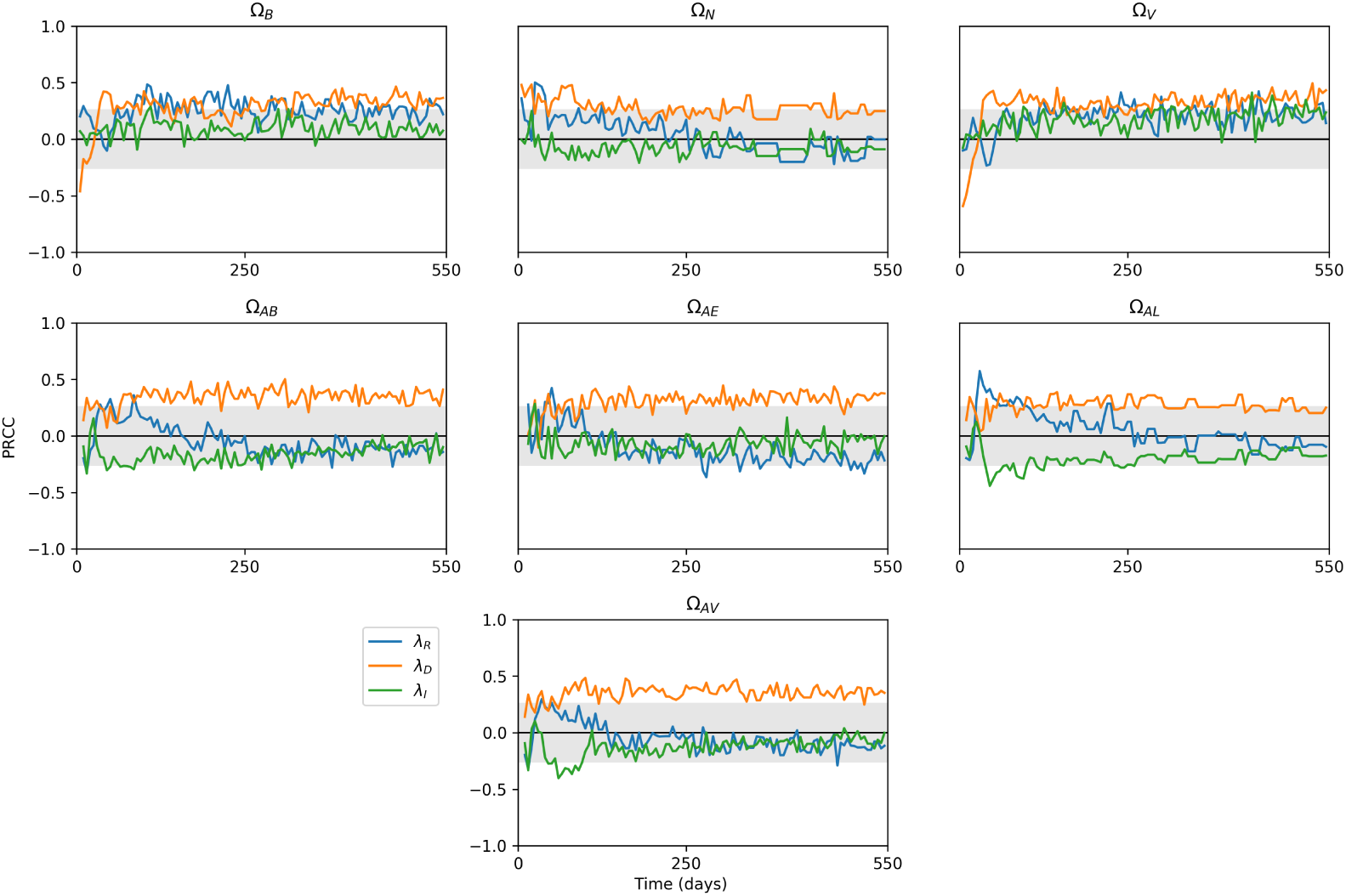
Sensitivity plots for bacterial replication parameters within *TBMetapopPy*, plotting the PRCC value of each parameter against one of the three outputs over time. Grey shaded area shows non-significance (p<0.01).

## 5. Discussion

Computational models of the within-host dynamics of TB have been used extensively to improve our understanding of how disease progresses within the body, particularly at the scale of a single lesion [29, 25, 53] and with some models looking at the disease over the whole lung and the associated lymph nodes [43, 44, 54, 55]. However, these models have not included the notions of heterogeneity of environmental conditions within the lung as shown in this paper, which are believed to be critical to the apical localisation of TB during the crucial post-primary stage [14]. In our first model [31], we presented the first *in silico* model of a pulmonary TB infection that included environmental heterogeneity and bacterial dissemination to show how the environmental conditions at the apex of the lung can favour bacterial growth and lead to larger bacterial loads at that location during latency. In this work, we have expanded on that initial model by including the whole life-cycle, from initial infection with bacteria to post-primary disease.

The results of our numerical simulations have shown the plausibility of environmental heterogeneity within the lung being the driving factor for an apical localisation in the organ of post-primary TB. Within our model, we have set the initial infection to begin in the lower regions of the lungs. As this progresses, the bacteria proliferate until their numbers are brought under control by the adaptive immune response. However, in order to trigger this adaptive response, antigen-presenting cells (dendritic cells and macrophages in our model) must traffic to the draining lymph nodes. By doing so, these cells present a means of bacterial dissemination, since any alive bacteria inside the cells are transferred along with the cell. This establishes a secondary infection within the lymphatic system. There is then a short window in our model whereby bacteria are present in the lymph nodes but before a solid granuloma has formed there, and it is during this period that the free bacteria are able to access the blood stream and re-seed the lung. Various smaller lesions form, but due to the adaptive immune response already being established, these lesions do not reach the CFU counts of the single lesion seen in the initial infection. These lesions are heterogeneous, with lesions towards the apex reaching a greater CFU count than those lower down. Once an external event reduces the immune response (by knocking out some of the recruitment of T cells), the numerous lesions present begin to expand, and it is in this region that the lesions reach the dangerous bacterial loads seen during primary infection. Our results also show that the composition of the lesions at different regions is heterogeneous, with apical lesions having a greater proportion of extracellular bacteria, especially those of a replicating phenotype. It would be interesting to make a comparison between the apical location identified in this paper with radiological lesions, but such a study was beyond the scope of this work.

Our sensitivity analysis shows that the skew of perfusion values has a significant effect on the total number of bacteria and the average lesion CFU during the all stages, whereas the skews of ventilation and drainage have limited impact. Therefore, we can interpret this as showing that the heterogeneity of perfusion may be the main environmental factor that contributes to the apical localisation, probably due to the reduced immune response it causes there, and this is shown by the strong sensitivity of Ω_*AV*_ to *S* _*Q*_. Differentials in blood flow have long thought to contribute to this apical localisation, with bed rest (which alters the scale and direction of blood flow due to the change in the host’s posture [56]) previously being shown to have positive effects on TB recovery [19] and this is the first modelling evidence to support this idea.

The results of our sensitivity analysis also highlight other mechanisms that may contribute to disease outcome. We have demonstrated the importance of T cells within the model, as the majority of T cell related parameters had significant influence on all three of our disease outputs. For our model, extracellular replication of dormant has a greater impact on disease outputs during the latency and post-primary stage than intracellular replication. Thus, it may be more important to focus treatments on destroying these extracellular bacteria than those in the intracellular niche to improve treatment.

Our *TBMetapopPy* environment models a single individual, with environmental attributes determined at the beginning of simulation remaining constant throughout. In reality, whilst the environment of the lung affects the disease, the reverse is also true, with the disease altering the anatomy of the lung, with tissue destruction and cavitation creating new routes of bacterial dissemination. Furthermore, our model simulates a single individual, with one set of environmental parameters being used for all simulations. However, no two human beings have the same lung, as lung morphology and physiology are affected by a number of factors, including height and body size [57], and sex [58, 59, 60]. Gender may be of particular interest to TB researchers, as there has been shown to be gender differences in responses to treatment: an analysis of the REMoxTB clinical trial revealed that men with cavitation showed poorer response to treatment than women with or without cavitation [61]. These differentials in disease outcome between genders may be caused by biological sex differences between male and female lung morphologies [58, 62], although the causes may also be socio-economic or cultural [63]. Further iterations of the model should also incorporate dynamic environments: as the disease progresses, it necessarily impacts the environment on which it occurs, through tissue damage and cavitation. Thus, differences between regions of the lung may become or less pronounced as the disease advances.

The topology utilised to model the lungs in *TBMetapopPy* is simplistic and yet is sufficient to show how environmental heterogeneities between different regions impact TB. In order to enhance the model and better reflect the real-world anatomy, different lung topologies could be used. The space-filling curve shown here is capable of filling any 2-dimensional shape and would extend to non-symmetrical shapes such as those found in the human lung. Furthermore, the lymphatic architecture of the model could be expanded, with a series of lymph nodes draining different regions of the lungs allowing for more complex transmission dynamics across the pulmonary environment.

During a TB infection, the interactions that occur between immune cells and bacteria are various and complex. In order to keep the model parsimonious, restrictions have been made on the number and type of immune cells included. Neutrophils play a major role in TB pathology, often being the first immune cells that encounter bacteria [64]. Separating neutrophils out into a separate compartment may allow for a more accurate representation of the real world dynamics, and to better understand cell-to-cell interactions and the impact the environment has on these, as neutrophils have strong interactions with macrophages [64]. Furthermore, the assumption of similar replication rates for dormant extracellular bacteria and intracellular bacteria may need further investigation, as intracellular bacteria have been shown to impact antigen-presentation, suggesting the bacteria are not entirely dormant [36, 65].

The focus of our model has been to simulate the disease in its untreated form. However, it is reasonable to assume that environmental heterogeneities also impact the pharmacokinetics of TB treatment, as all drugs will enter the environment via the blood and thus perfusion differentials may result in inadequate concentrations of chemotherapy reaching the areas of worst bacterial growth. The pharmacodynamics of the different drugs that are currently used to fight TB are affected by different environments and so a variety of different drugs could be included in the model that perform different actions dependent on their location. The ability to combine a whole-organ model of various lung morphologies with a range of possible treatment regimens would confer the ability to create ‘virtual clinical trials’, where new regimens could be trialled on synthetic patients first, allowing us to make predictions on their likely efficacy and thus better prioritise the regimens to be put into actual trials.

## Appendix A. Parameter estimation

The parameters used in the event dynamics, environment construction and initial conditions have been derived from existing literature where possible or estimated. Each parameter is given a baseline value for the purposes of running individual simulations (see Section 3), as well as a range, representing the biological uncertainty of the parameter, which is used for sensitivity analysis (see Section 4). As there are not many experimental or clinical studies focused on humans, many parameters are derived from animal models or have been estimated. Where parameters have been estimated, large ranges have been applied as well as a uniform distribution [66]. The uncertainty created by these estimated parameters is explored in Section 4.

### Environmental parameters

In the human body, blood flow to the base of the lungs is approximately 3 times that at the apex of the lungs, whilst ventilation at the base is approximately twice that the apex [22]. We use these observations to determine ranges of values for *S* _*V*_ and *S* _*Q*_ (see Table 3), and it is these values ultimately determine the amount of blood and air present in each section of the lung.

### Bacterial state changes

The oxygen values within each patch are relative values indicating the difference between ventilation and perfusion - thus, we assume a value of 1 for oxygen implies a balance between oxygen entering that area of the lung and oxygen being removed through oxygen exchange. Values above 1 indicate a surplus of oxygen and values below 1 indicate a deficit. We thus choose 1 as a half-sat value for bacterial state changes from replicating to dormancy and vice versa (*θ*_*ξ*_).

### Recruitment and death

Macrophage recruitment rates are based on known macrophage numbers within the lung [46] and determined based on these and the macrophage death rate taken from other experimental models [30]. For dendritic cell recruitment, *α*_*D*_, we assume that the dendritic population is approximately 10% of the macrophage population [43], and thus choose scaled values based on the macrophage recruitment rate derived from the literature. Similarly, enhanced dendritic cell recruitment, *θ*_β*D*_, is scaled based on the corresponding value for macrophages, *θ*_β*ML*_.

### Cell-to-cell interactions

Little experimental data exists upon which to derive accurate values for the rates of contacts and half-saturation values between specific cells during infection. Where possible, values have been derived from existing models. In cases where there is no data to draw on, the values have been assigned wide parameter ranges for the Sensitivity Analysis (see Section 4). For T cell destruction of bacteria, previous models have either destroyed all bacteria [29] or released a portion of them [44]. As such, we have chosen to use a baseline value of 0.5 for *ψ*_*κM*_ to signify half of all bacteria being destroyed, with a range that varies up to 1 (signifying all bacteria being destroyed).

