## Supplementary Materials for "Modelling the effects of environmental heterogeneity within the lung on the tuberculosis life-cycle"

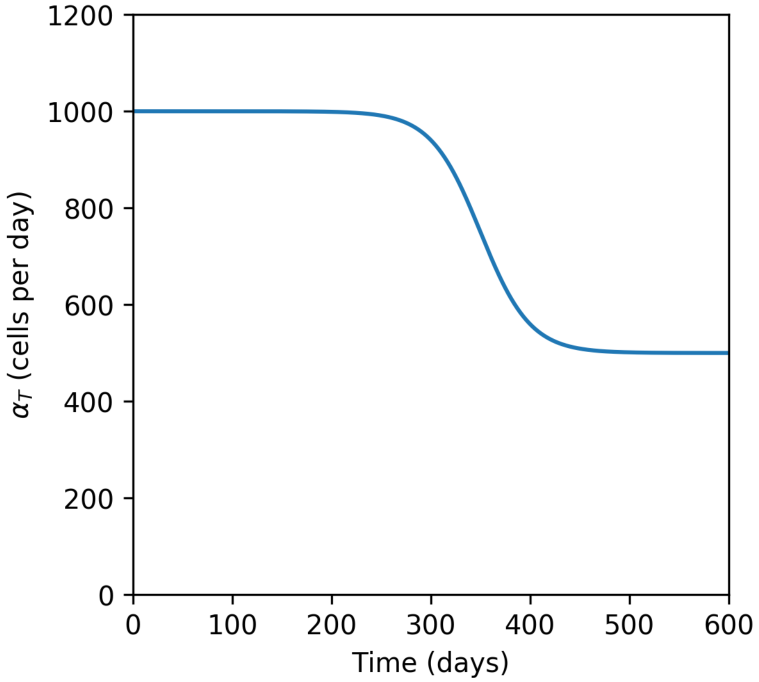


Figure S1: Change in T cell recruitment rate ($\alpha_T$) over time, when $\alpha_T(0) = 1000$

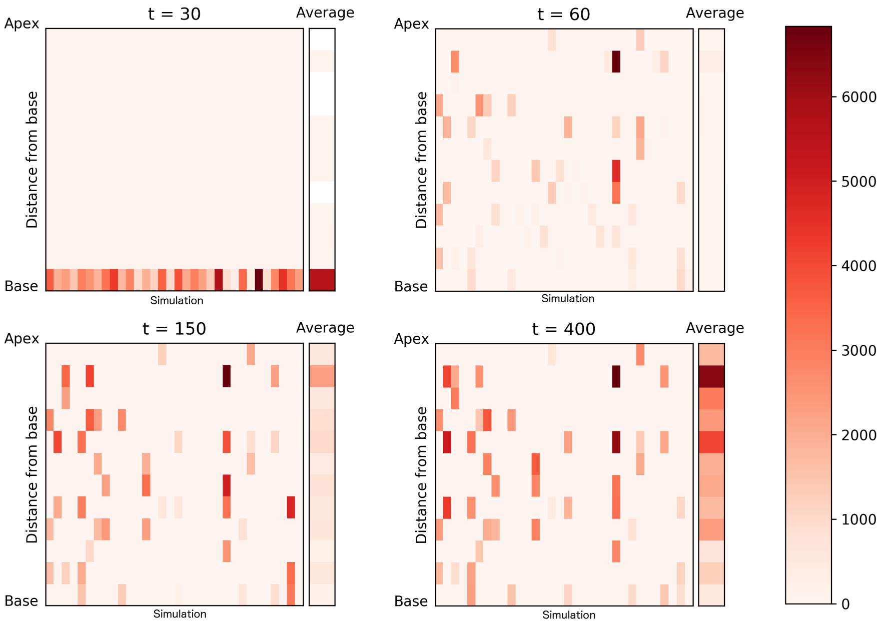


Figure S2: Average lesion size (number of bacteria) at each of 12 evenly sized horizontal slices of lung at different time-steps during the simulation. Columns represent individual simulations, with the final column representing the average across simulations.

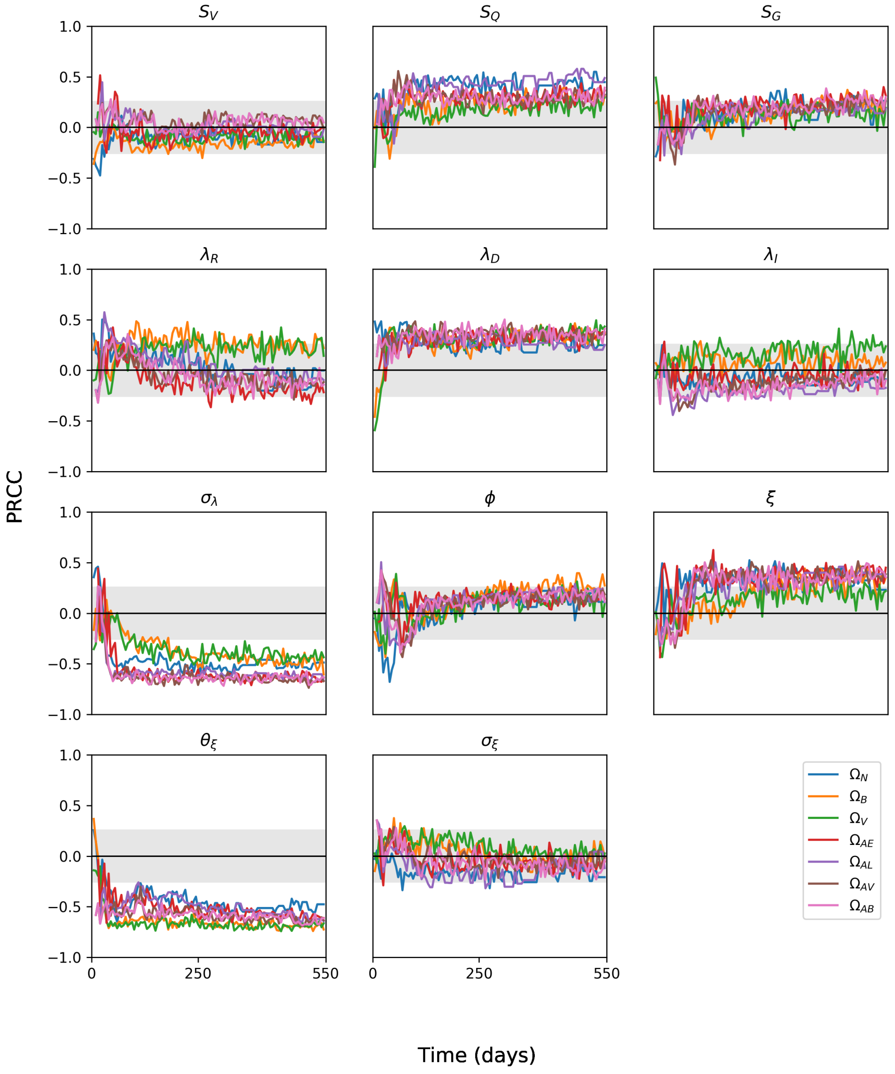


Figure S3: Sensitivity plots for various parameters within *TBMetapopPy*, plotting the PRCC value of the parameter against the model outputs over time. Grey shaded area shows non-significance (p<0.01).

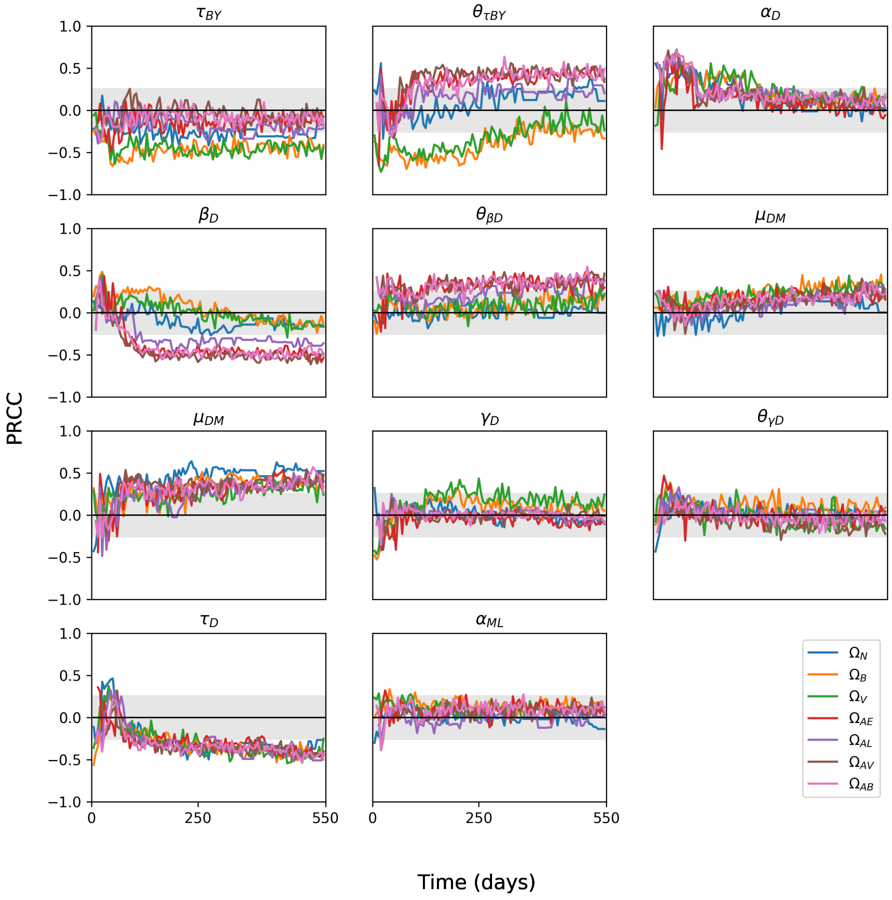


Figure S4: Sensitivity plots for various parameters within *TBMetapopPy*, plotting the PRCC value of the parameter against the model outputs over time. Grey shaded area shows non-significance ($p<0.01$).

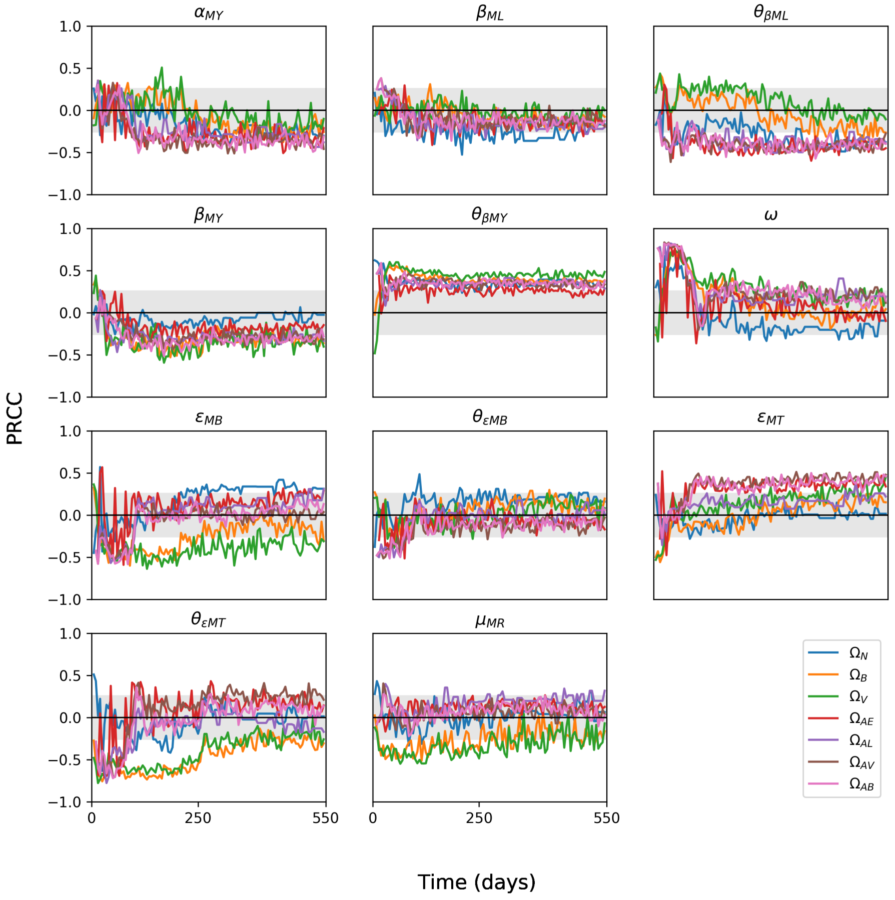


Figure S5: Sensitivity plots for various parameters within *TBMetapopPy*, plotting the PRCC value of the parameter against the model outputs over time. Grey shaded area shows non-significance (p<0.01).

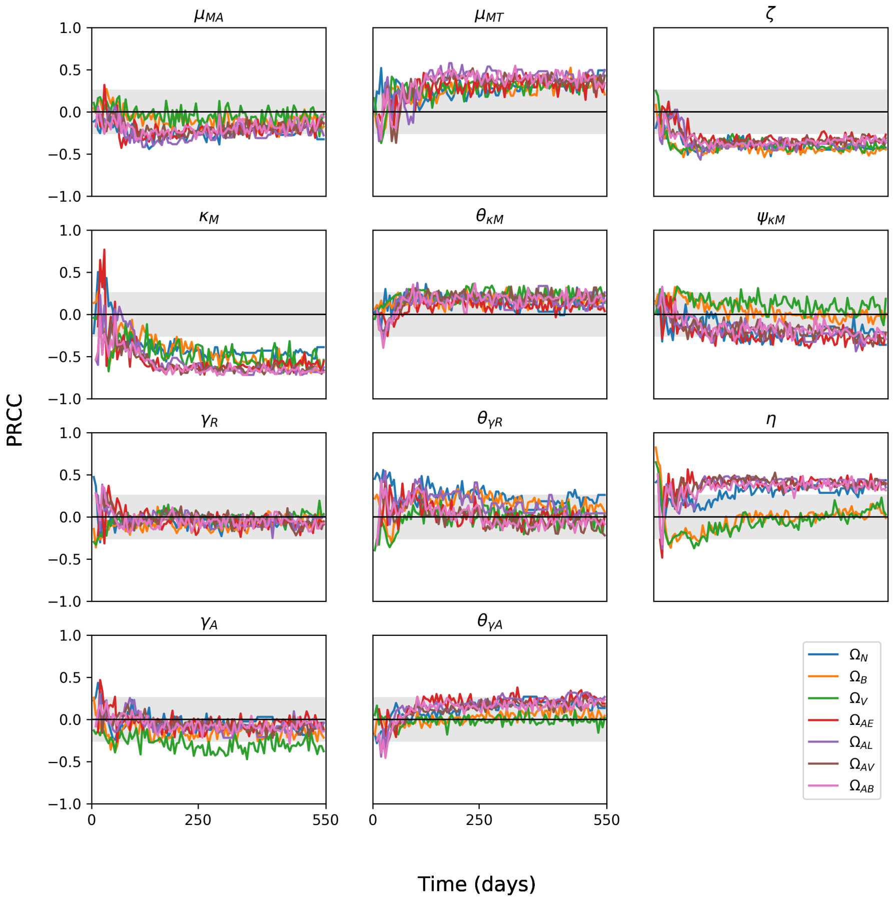


Figure S6: Sensitivity plots for various parameters within *TBMetapopPy*, plotting the PRCC value of the parameter against the model outputs over time. Grey shaded area shows non-significance (p<0.01).

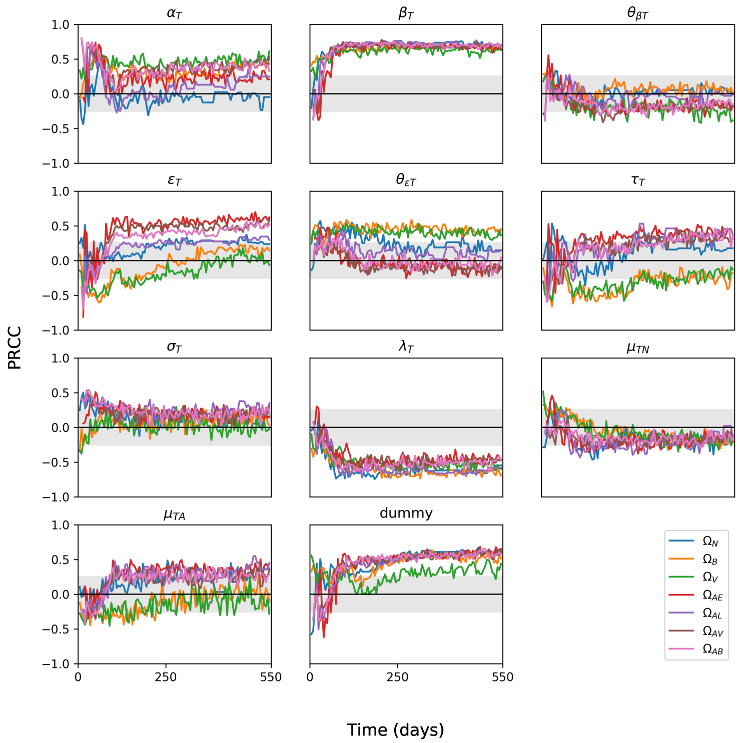


Figure S7: Sensitivity plots for various parameters within *TBMetapopPy*, plotting the PRCC value of the parameter against the model outputs over time. Grey shaded area shows non-significance (p<0.01).
